# De Novo Negatively Charged Binders Targeting MMLVRT Nucleic Acid Binding Sites Overcome Stability-Activity Trade-offs

**DOI:** 10.1101/2025.03.15.643416

**Authors:** Yibo Zhu, Hongwei Liu, Fang Qu, Yingzhi Wang, JuanJuan Yao, Shumin You, Lin Hua, Chunlei Ge, Hong Yao, Tao Li

## Abstract

The limited thermostability and storage-induced inactivation of Moloney murine leukemia virus reverse transcriptase (MMLV RT) have constrained its applications. In this study, a high-stability mutant, MMLV RT-SV, was generated through a multi-site mutagenesis strategy targeting the nucleic acid-binding pocket. Through de novo protein design, three highly negatively charged binders with affinities of 45 nM, 362 nM, and 829 nM were developed to specifically target the positively charged nucleic acid-binding region on the surface of MMLV RT-SV. Experimental results demonstrated that these binders formed complexes with the enzyme via electrostatic interaction, significantly enhancing the thermostability and long-term storage stability of MMLV RT-SV while not affecting its RNA-dependent DNA polymerase function. This work pioneers the rational design of negatively charged binders targeting strongly positively charged nucleic acid-binding sites, overcoming the traditional stability-activity trade-off inherent in site-directed mutagenesis and directed evolution. The proposed strategy not only provides an innovative solution to address the thermal sensitivity and storage instability of reverse transcriptase but also establishes a novel paradigm for De Novo-based precision engineering of enzyme functions, demonstrating significant potential in molecular diagnostics, gene editing, and related fields.

## 1. Introduction

Moloney Murine Leukemia Virus Reverse Transcriptase (MMLV RT), a monomeric protein extracted from the virus with a molecular weight of 75 kDa, has a structure comprising the finger domain, palm domain, thumb domain, connection domain, and RNase H domain, which are responsible for nucleic acid binding, catalyzing DNA synthesis, maintaining enzyme conformation, and cutting RNA-DNA hybrid chains, respectively[1]. This enzyme possesses RNA-dependent DNA polymerase activity, RNase H activity, and terminal transferase activity that adds nucleotides to the DNA chain end in a template-independent manner [2]. It is widely used in gene cloning and in vitro amplification and has extensive applications in bioengineering, biomedical, and molecular biological fields, including cDNA synthesis, RT-PCR, gene cloning, gene expression analysis, constructing gene libraries, gene editing, genome sequencing, and viral diagnostics [3,4]. During the COVID-19 pandemic, MMLV RT was widely used for SARS-CoV-2 RT-PCR detection, highlighting its importance in clinical diagnostics [5,6]. However, MMLV RT has low thermostability and is typically used at 37℃. The secondary structure of RNA and high GC content templates can hinder primer binding and reverse transcription at this temperature, leading to reduced cDNA synthesis efficiency and limiting its application under high-temperature conditions. Moreover, long-term storage and repeated freeze-thaw cycles can significantly affect the activity and thermostability of reverse transcriptase, thereby adversely affecting experimental results and practical applications.

To address these issues, researchers have attempted to enhance the thermostability of MMLV RT to enable it to work at higher temperatures, thereby reducing the impact of RNA secondary structures and improving reverse transcription efficiency. Currently, strategies to improve MMLV RT stability mainly rely on directed evolution or site-directed mutagenesis [7,8]. For instance, to overcome the deficiency of low thermostability, researchers have significantly enhanced the enzyme’s activity and half-life at high temperatures (50-60℃) through multiple mutant combinations (such as enhancing structural rigidity) and optimizing the template-primer binding interface [9,10]. However, single-site mutations may not comprehensively enhance enzyme stability, and the design of site-directed mutagenesis may be limited in the absence of complete structural information. Additionally, the efficiency of directed evolution is limited by structural information, the scale of mutant libraries, and the effectiveness of screening methods. It may also lead to a trade-off between enzyme activity and stability, meaning that enhancing stability may sacrifice activity, and vice versa. Thus, different approaches are essential to improve the heat stability and long-term storage capacity.

Nucleic acid aptamers and small molecule auxiliary reagents stabilize proteins through “structural complementary binding” and “physical chemical interactions”, respectively, and have significant application value in scenarios with high specificity requirements and large-scale applications [11,12]. Moreover, protein-protein interactions (PPI) are widely used to enhance enzyme stability. For example, specific antibodies, molecular chaperones, or inhibitors can form stable complexes with enzymes, effectively enhancing their thermostability, resistance to degradation, and long-term storage performance[13–15]. This indicates that the development of new enzyme-binding proteins has the potential to overcome the limitations of MMLV RT poor thermostability and activity reduction caused by long-term storage and repeated freeze-thaw cycles without affecting enzyme function.

In recent years, with the rapid development of artificial intelligence technology, deep learning-based protein de novo design technologies, such as AlphaFold3, RoseTTAFold, RFdiffusion and ProteinMPNN, have achieved revolutionary progress in the field of protein engineering[16–18]. These technologies have provided a new paradigm for developing high-affinity and high-specificity functional protein binders by accurately predicting protein three-dimensional structures and interaction interfaces[19,20]. In particular, they have shown great potential in targeted enzyme activity regulation, making it possible to directly target protein design binders to enhance stability. By accurately predicting protein three-dimensional structures and interaction interfaces, researchers can design functional protein binders with high affinity and specificity, thereby directly targeting proteins to enhance their stability. This not only has significant theoretical importance but also demonstrates great practical value.

In this study, we found that the multi-site stable variant (MMLV RT- SV) targeting the nucleic acid-binding pocket of MMLV RT, compared to the wild-type MMLVRT (MMLV RT-WT), showed significantly enhanced stability after binding to nucleic acid complexes based on templates/primers. Furthermore, using the high-stability mutant MMLV RT-SV as the core, we designed and screened a series of protein binders that can specifically bind to the strongly positively charged nucleic acid-binding pocket of MMLV RT, based on deep learning-driven protein de novo design technology. We obtained three strong binders with affinities of 45 nM, 362 nM, and 829 nM, respectively. These three strong binders significantly improved the thermostability and activity of MMLV RT-SV under long-term storage conditions. In addition, we found that the binding of the binder to MMLV RT-SV does not affect the normal transcription function of MMLV RT-SV. This innovative approach of combining multi-site mutations with de novo designed binders to enhance MMLV RT-SV stability provides a solution and possibility for the important application of MMLV RT-SV in bioengineering, biomedical, and molecular biological fields. It also demonstrates the possibility of designing strongly negatively charged binders for strongly positively charged nucleic acid-binding sites for the first time. This study not only provides an innovative solution to the technical bottleneck of poor thermostability and reduced enzyme efficiency due to long-term storage in MMLVRT applications but also opens up a new research direction for the optimization and precise application of in vitro diagnostic technologies.

## 2. Results

### 2.1 Engineered MMLV RT Variant Enhances Thermostability and Catalytic Efficiency via Nucleic Acid-Binding Pocket Optimization

The stable binding of Moloney Murine Leukemia Virus Reverse Transcriptase (MMLV RT) to template/primer (T/P)-centered nucleic acid complexes is a critical determinant for its efficient catalysis of RNA-dependent DNA synthesis in reverse transcription polymerase chain reaction (RT-PCR) and in vitro diagnostic technologies. Given the strong negative charge characteristics of T/P nucleic acid complexes, we developed a multi-site stabilized variant (MMLVRT-SV: D584N E70K E303R T307K W314F L436G N455K) targeting the nucleic acid-binding pocket of MMLV RT, based on improved template binding and optimize to thermal denaturation (Figure 1A). Compared to wild-type MMLV RT (MMLVRT-WT), MMLVRT-SV incorporates crucial positive charge- enhancing and stability-improving mutations in the nucleic acid-binding pocket (Figure 1B). The RMSD of 0.422Å between their AlphaFold3- predicted structures suggests these mutations preserve proper folding and maintain normal protein expression (Figure 1C).

**Figure 1.**
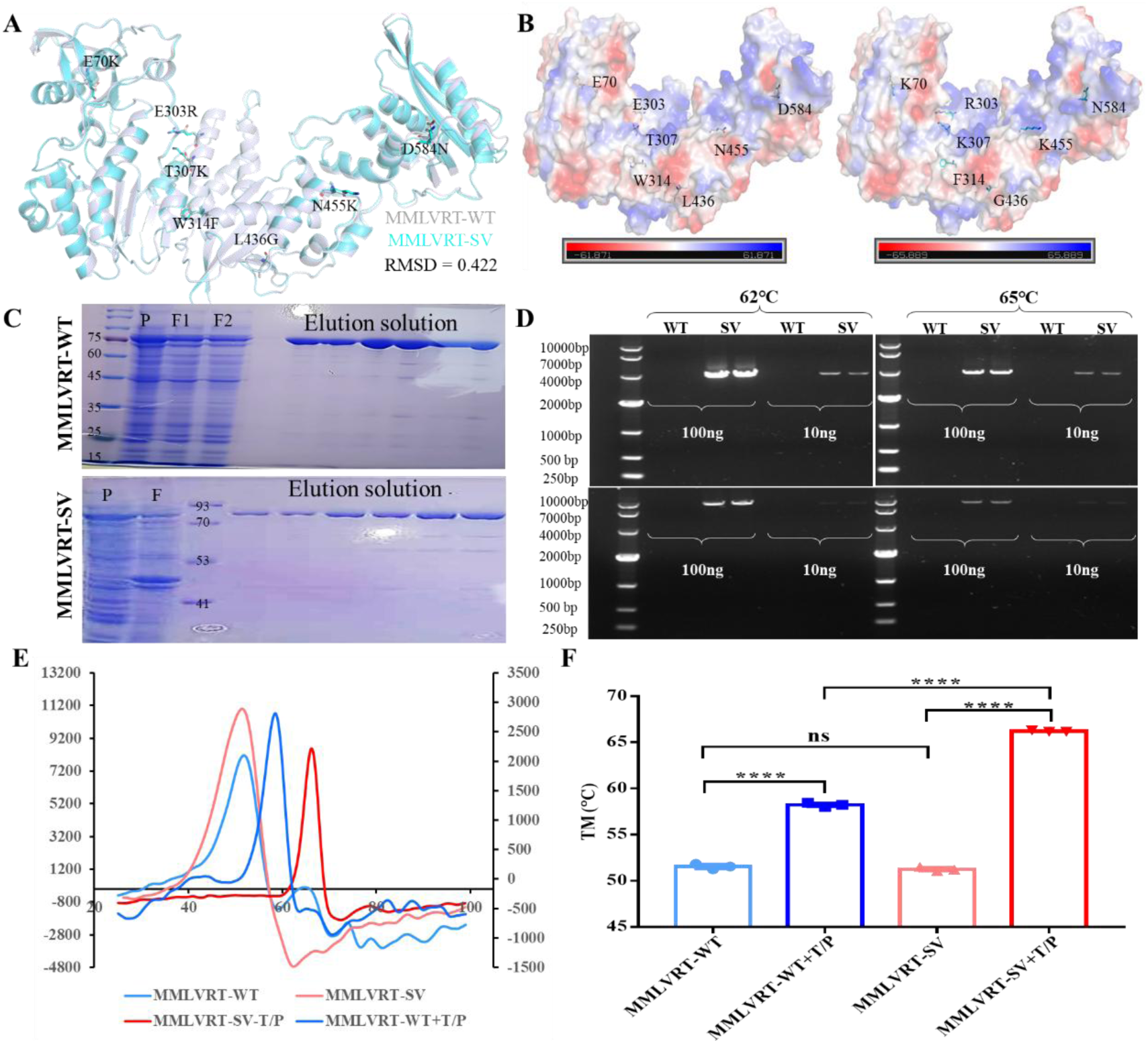
MMLVRT-SV increased the positive charge in the nucleic acid - binding pocket and stable site mutations, enhancing the thermal stability of T/P binding. (A) Display of MMLVRT-SV mutation sites, with the MMLVRT-SV structure shown as a cyan cartoon and MMLVRT-WT as a gray cartoon. (B) Surface charge distribution maps of MMLVRT-WT (Left) and MMLVRT-SV (Right). (C) PAGE gel results of protein expression and purification for MMLVRT-WT (top) and MMLVRT-SV (bottom). (D) Agarose gel identification of reverse transcription results of MMLVRT- WT and MMLVRT-SV at different temperatures (62℃ and 65℃) and with different molecular weight templates (4K and 10K). The amount of MMLV added was 250 ng. The final concentrations of 293T mRNA in the 20 μl system were 5 ng/μl and 0.5 ng/μl. (E-F) DSF identification of melting curves (E) and melting temperature statistical chart (F) of MMLVRT-WT and MMLVRT-SV alone and in combination with T/P. 500ng MMLV RT-WT or 500ng MMLV RT-SV, 2.5µM of the T/P nucleic acid complex in DSF system.

Notably, MMLVRT-WT exhibited a drastic reduction in transcriptional efficiency as temperatures increased, losing virtually all activity at 50-55°C (Supplementary Figure 1). In stark contrast, the multi- site engineered variant MMLVRT-SV demonstrated superior thermal resilience, maintaining stable reverse transcription activity even at temperatures >60°C across diverse template sizes (Figure 1D). Under these high-temperature conditions, MMLVRT-SV consistently produced high- yield products in RT-PCR (Figure 1D), whereas MMLVRT-WT failed to generate detectable products (Figure 1D & Supplementary Figure 1). This pronounced divergence in performance directly confirms that multi-site engineering of the nucleic acid-binding pocket significantly enhances both sensitivity and stability during reverse transcription.

Differential scanning fluorimetry (DSF) assays revealed that MMLVRT-SV and MMLVRT-WT shared comparable intrinsic thermal stability (Tm=51.53°C vs 51.25°C, Figure 1E&F). However, upon T/P complex binding, MMLVRT-SV showed markedly enhanced complex stabilization (ΔTm=15°C vs 7°C for WT), achieving an 8°C higher Tm (66.20°C vs 58.19°C, Figure 1E&F). These results demonstrate that MMLVRT-SV optimized nucleic acid binding significantly improves reaction specificity and adaptability through enhanced complex stabilization. The variant elevated optimal temperature substantially boosts cDNA synthesis efficiency and quality, indicating superior RT performance with important implications for molecular biology research and diagnostic technology advancement.

### 2.2 Enhancing MMLV RT Stability: A Deep-Learning-Driven De Novo Binder Design Approach by Targeting Nucleic Acid-Binding Pocket

To address the decline in enzymatic activity and thermal stability of Moloney murine leukemia virus reverse transcriptase (MMLV RT) during long-term storage or repeated freeze-thaw cycles, this study focused on the topological features of the nucleic acid - binding pocket of MMLV RT and employed a deep-learning-driven de novo protein design strategy to develop binders targeting this region.

The study commenced by predicting the structure of the MMLV RT- SV nucleic acid complex using AlphaFold3 (Figure 2A). Key hotspot residues at the binding interface were identified via PDBePISA and used as constraints in the RFdiffusion platform for geometrically constrained pocket topology matching design (Figure 2B, Supplementary Table 1). Diffusion models were utilized to generate α-helical scaffolds, and 12 high-matching scaffolds were selected based on interface contact density and shape complementarity (Supplementary Figure 2). For each selected scaffold, sequence-space exploration was conducted using ProteinMPNN, generating five sequences per scaffold to form a candidate binder library. These candidates underwent dual validation through AlphaFold3 for binder-apo structure prediction and MMLV RT-binder complex docking. Based on interface pTM (ipTM) and predicted local distance difference test (pLDDT) score, as well as the overlay of apo and complex structures, 32 candidates meeting both geometric and energy criteria were identified.

**Figure 2.**
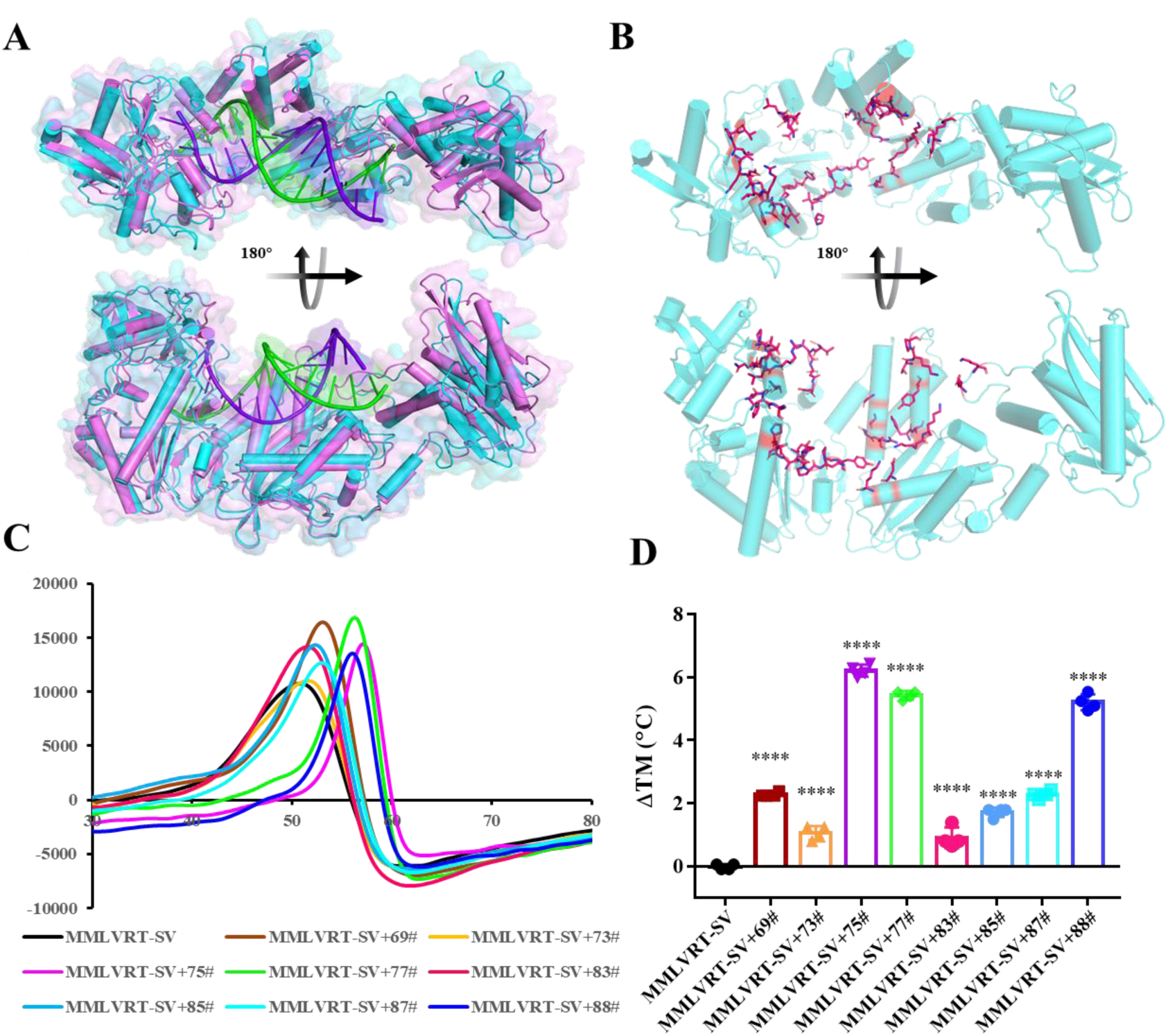
Hotspot selection in MMLVRT-SV and de novo binder design and screening. (A) AlphaFold3 predicted the structure of the MMLVRT - SV - T/P complex, with the MMLVRT-SV monomer structure (apo) displayed as a pink cartoon, and the MMLVRT- SV-T/P complex shown as a rose cartoon. The template RNA is represented as a green cartoon, and the primer DNA is depicted as a purple cartoon. (B) PDBePISA analyzed the key sites in the interaction interface of the MMLVRT-SV-T/P complex and their positions in the pocket are shown. (C-D) DSF analysis was performed to determine the melting curves and shifts in melting temperature (ΔTm) of 0.74 μM MMLVRT-SV following the addition of 4.4 μM candidate binder in the final system.

The candidate sequences were synthesized into plasmids for prokaryotic expression, resulting in the successful production of eight binder proteins (Supplementary Figure 3). Differential Scanning Fluorimetry (DSF) analysis revealed that binders 75#, 77#, and 88# significantly increased the melting temperature (ΔTm = 6.2°C, 5.4°C, and 5.2°C, respectively) of MMLV RT-SV (Figure 2C & D). This enhancement in thermal stability aligns with the design model, where the binder “structurally clamps” the catalytic core. This study is the first to demonstrate that computationally designed closure of the nucleic acid - binding domain can effectively improve reverse transcriptase stability, offering a new paradigm for enzyme engineering.

### 2.3 Candidate Binders Bind to MMLV RT-SV and Significantly Enhance Its Thermal Stability

To better confirm whether the three binders enhance the stability of MMLV RT-SV in a concentration-dependent manner and to clarify their binding affinity. We initially investigated the thermal stability enhancement of MMLV RT-SV by the three binders under different concentration conditions. The results showed that the Tm value of MMLV RT-SV increased significantly with the increasing concentration of the binder, indicating that the binding of the binder to MMLV RT-SV increased its thermal stability in a concentration - dependent manner (Figure 3A). The concentrations corresponding to the half-maximal ΔTm for Binder 75#, 77#, and 88# were 1.52±0.42 µM, 0.9±0.13 µM, and 0.72±0.09 µM, respectively (Supplementary Figure 4). This indicated that Binder 75# was the most effective in enhancing the thermal stability of MMLV RT-SV, followed by Binder 77#, and Binder 88# was relatively less effective. In terms of the maximum temperature increase after the addition of the three binders, the values were 9.108±1.496°C, 6.221±0.48°C, and 5.595±0.334°C for Binder 75#, 77#, and 88#, respectively, with Binder 75# showing a greater maximum temperature increase (Supplementary Figure 4).

**Figure 3:**
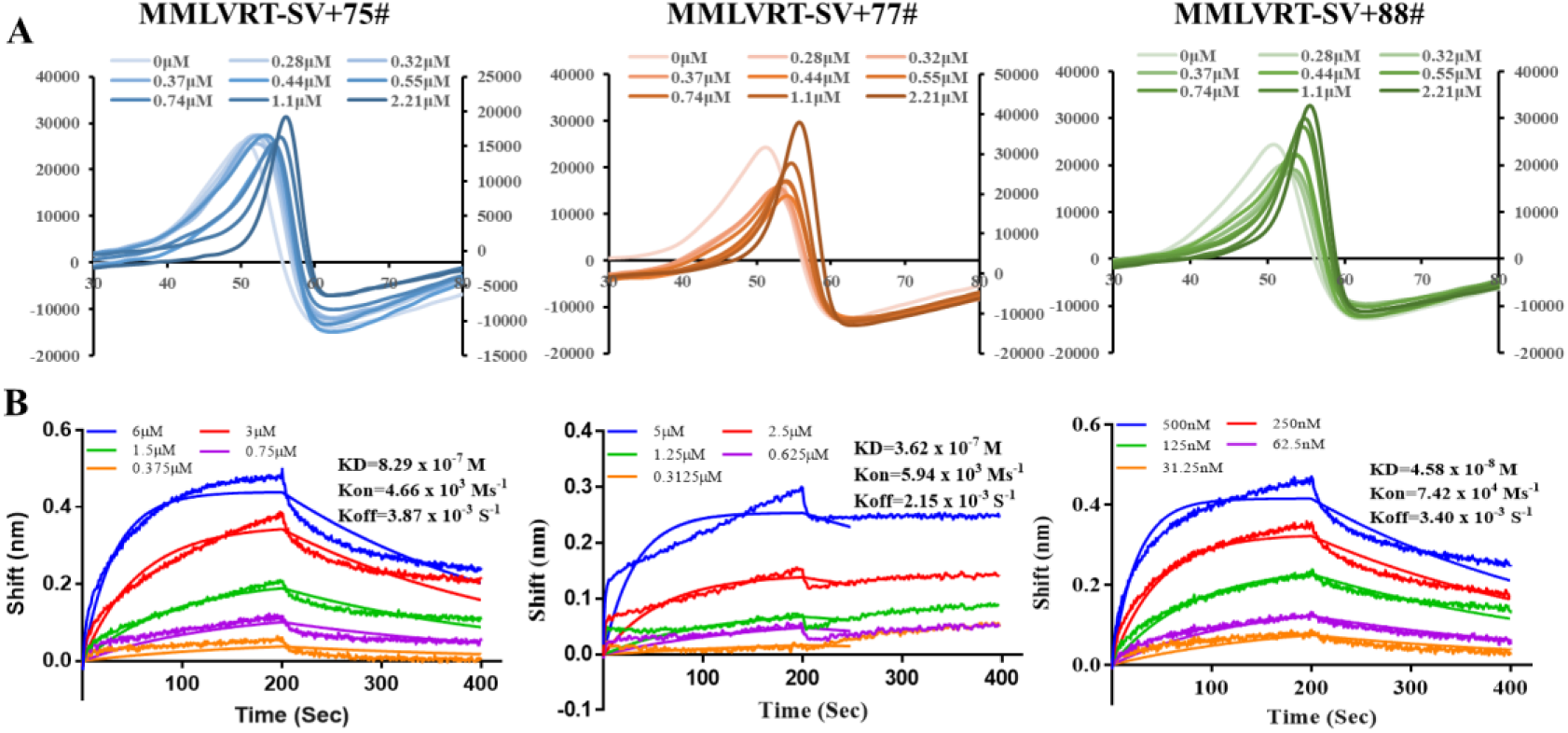
Affinity determination of Binder75#, 77# and 88# for MMLV RT-SV and their effect on enhancing the thermal stability of MMLV RT-SV. (A) DSF determined changes in the thermal denaturation curves of MMLV RT-SV under the condition of gradient concentrations (0 to 2.21μM) of Binder added. (B) DSF determined the change in melting temperature values (ΔTM) of MMLV RT-SV under the condition of gradient concentrations (0 to 2.21μM) of binders added. (D) Bio-Layer Interferometry (BLI) determined the affinity of Binder75#,77# and 88# for MMLV RT-SV.

Subsequently, we used Bio-Layer Interferometry (BLI) to determine the binding affinities of the three binders with MMLV RT-SV. The results showed that the binding affinities of Binder 75#, 77#, and 88# for MMLV RT-SV were 829 nM, 362 nM, and 45.8 nM, respectively (Figure 3B). These data indicated that we successfully designed and screened binders with nanomolar-level binding to MMLV RT-SV as intended, and these binders enhanced the thermal stability of MMLV RT-SV in a concentration-dependent manner.

### 2.4 Candidate Binders Enhance MMLV RT-SV Stability Without Affecting Reverse Transcription Reaction

To verify whether the binding of the three candidate binders to MMLV RT-SV would affect the ability of MMLV RT-SV to bind to T/P (template/primer), thereby impacting the reverse transcription process and cDNA synthesis, we investigated the reverse transcription and cDNA synthesis in the MMLV RT-SV-binder complex system after adding T/P. The results showed that the binding of MMLV RT-SV-binder did not affect the reverse transcription process and cDNA synthesis of MMLV RT-SV (Figure 4A).

**Figure 4:**
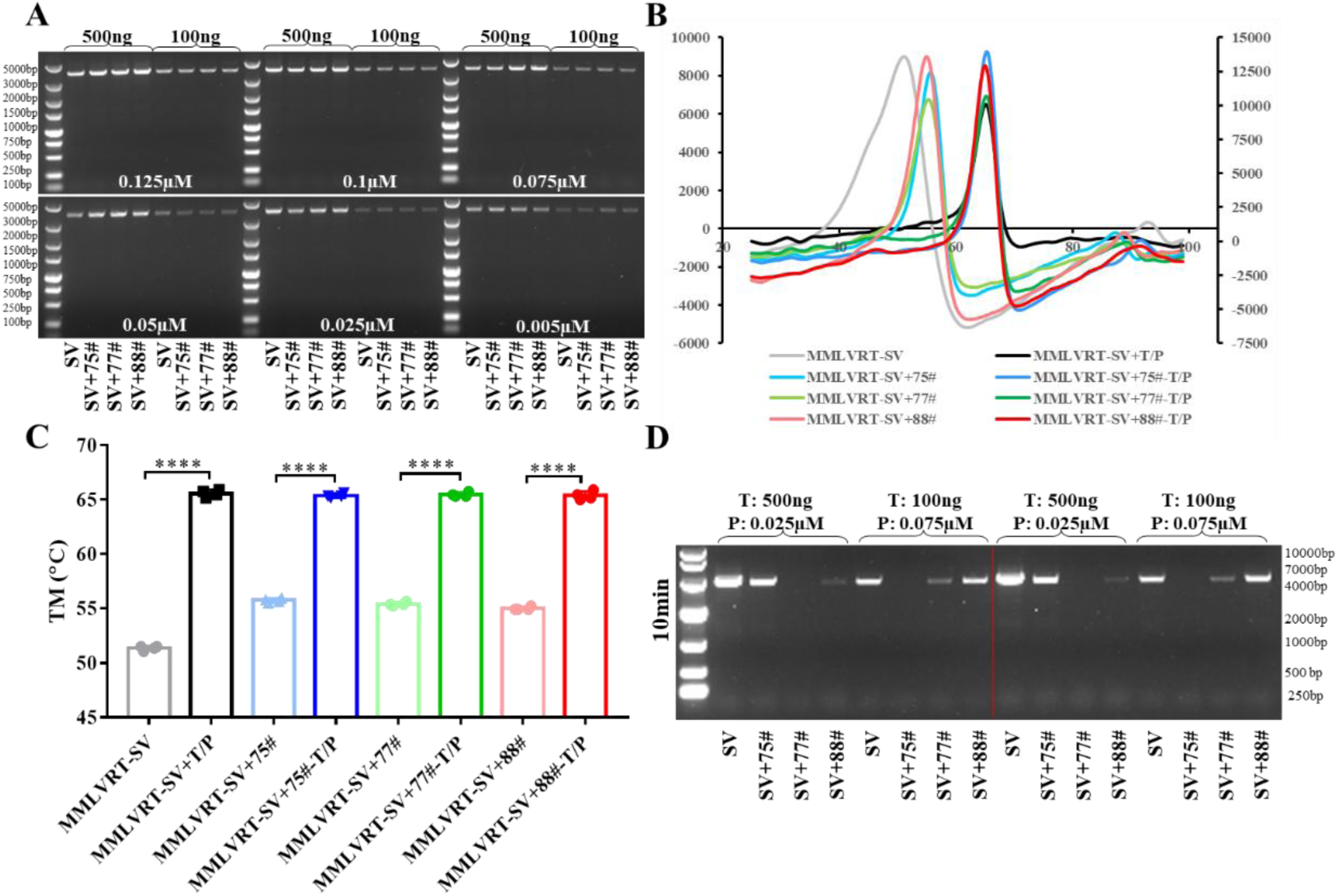
Binder75#, 77# and 88# addition and their impact on the reverse transcription process and cDNA synthesis by MMLV RT-SV. (A) After pre-mixing 7.36 µM Binder75#, 77#, and 88# with 250 ng MMLV RT-SV, at 16°C for 30 min, the mixture was left overnight. Subsequently, other components, including dNTP Mix, Oligo dT (18T) Primer (final concentration 0.125-0.005µM), and RNase Inhibitor, were added, and the reaction was performed at 37°C for 30 min.The reverse transcription process was carried out. With 50uM MMLV RT-SV+Binder Mix, 1ug template, and 10μM primer, reverse transcription was performed at 37℃, and the cDNA products were identified through agarose gel. (B&C) DSF determined melting curves (B) and statistical results of melting temperature (C) of the pre-mixed solution of 2.2µM Binder75#, 77# and 88# with 0.368 µM MMLV RT-SV, with or without the addition of 2.5µM T/P. (D) The reaction time was shortened to 10 min, and reverse transcription was carried out under the conditions of Primer: 0.025µM/ Template:500ng or Primer: 0.075 µM/Template:100ng, with the cDNA products identified through agarose gel.

To further explore the reason, we conducted DSF experiments again to investigate the impact of adding T/P on the thermal stability of the MMLV RT-SV-Binder system under binder -bound conditions. The experimental results showed that, consistent with the previous results, the addition of Binder significantly enhanced the thermal stability of MMLV RT-SV (Figure 4B & C). Surprisingly, when T/P was added, the melting temperature of the MMLV RT-SV-Binder-T/P system was the same as that of the MMLV RT-SV-T/P system with T/P added alone, both at 65℃ (Figure 4B & C). Moreover, in the DSF experiments of the binder alone, T/P alone, and the Binder-T/P mixture, no significant signal values were detected, and the melting temperatures could not be evaluated (Supplementary Figure 5). This indicates that when T/P is present, the binder in the MMLV RT-SV-Binder system may be displaced by T/P through competition, and the binder did not bind to other sites of MMLV RT-SV. This further explains why the binding of the binder to MMLV RT- SV does not affect the reverse transcription process and cDNA synthesis of MMLV RT-SV, that is, the addition of T/P competitively replaced the binding of the Binder to MMLV RT-SV. It also indirectly confirmed that our designed Binder targeted the nucleic acid -binding site of MMLV RT- SV, and the binding ability of T/P to MMLV RT-SV was significantly stronger than that of the Binder to MMLV RT-SV (Figure 4B & C).

To investigate the competitive binding mechanism between Binders and the template/primer (T/P) complex in MMLV RT-SV, we systematically modulated T/P ratios (500 ng/0.025 μM vs. 100 ng/0.075 μM) and reduced reverse transcription time (Figure 4D). Under standard 30-minute reactions, cDNA synthesis remained unaffected, whereas shortening the reaction time to 10 minutes revealed distinct inhibitory effects of Binders on reverse transcription. Notably, dose-dependent suppression patterns varied significantly between conditions: In the high- template system (500 ng/0.025 μM), Binder77# completely abolished cDNA synthesis, Binder88# showed partial inhibition, and Binder75# exhibited minimal interference. Conversely, in the low-template system (100 ng/0.075μM), Binder75# induced complete suppression, Binder77# partially inhibited synthesis, and Binder88# showed negligible effects (Figure 4D). These findings confirm a concentration-dependent, competitive binding mechanism and highlight the critical role of Binder chemistry in modulating reverse transcriptase activity, providing a foundation for rational design of enzyme-specific inhibitors.

In general, these findings not only confirmed that the Binder successfully targeted the nucleic acid-binding pocket but also enhanced the thermal stability of MMLV RT-SV without affecting its reverse transcription and cDNA synthesis functions and activities. This is of great significance for applications in molecular diagnosis, gene editing, and other fields.

### 2.5 Do Novo designed binders significantly enhance the thermal stability and long-term storage performance of MMLV RT-SV

To further explore the impact of binder binding on MMLV RT-SV resistance to performance decay caused by long-term storage, we conducted an enzyme activity accelerated thermal stability experiment at 37℃. In this experiment, the enzyme was placed at 37℃ for different numbers of days, and RT-qPCR was performed on equal amounts of the enzyme at fixed time points to investigate the changes in its performance decay. We investigated the changes in CT values in qPCR after reverse transcription of equal amounts of mRNA by different MMLV RT-SV after 10 days of storage. The results showed that the CT value of the MMLV RT- SV group gradually increased with the extension of storage time (Figure 5A), indicating that the efficiency of MMLV RT-SV in reverse transcription of equal amounts of mRNA templates gradually decreased with the increase in the number of days in the accelerated thermal stability experiment. In contrast, the CT values of the MMLV RT-SV-binder group fluctuated less, especially in the MMLV RT-SV-77# Binder group. Even with the increase in the number of days in the accelerated thermal stability experiment, its CT value remained stable at the same level as day 0, and the overall CT fluctuation was maintained within 0.65 cycles (with a maximum of 1.66 cycles) (Figure 5A & B). The 75# binder was next, with the overall CT fluctuation maintained within 0.38 cycles (with a maximum of 2.175 cycles), and the 88# binder was relatively worse, with a maximum CT fluctuation of 4.52 cycles and an average of 2.3 cycles. Furthermore, compared with the day 0 and day 10 groups, the ΔCT value of the MMLV RT-SV group differed by nearly 8 cycles, while the ΔCT values of the 75# and 77# binder groups remained almost unchanged, and the CT value of the 88# group differed by nearly 4 cycles (Figure 5C & D). These data indicate that the designed binders not only enhance the thermal stability of MMLV RT-SV but also maintain the long-term storage stability of the enzyme performance.

**Figure 5.**
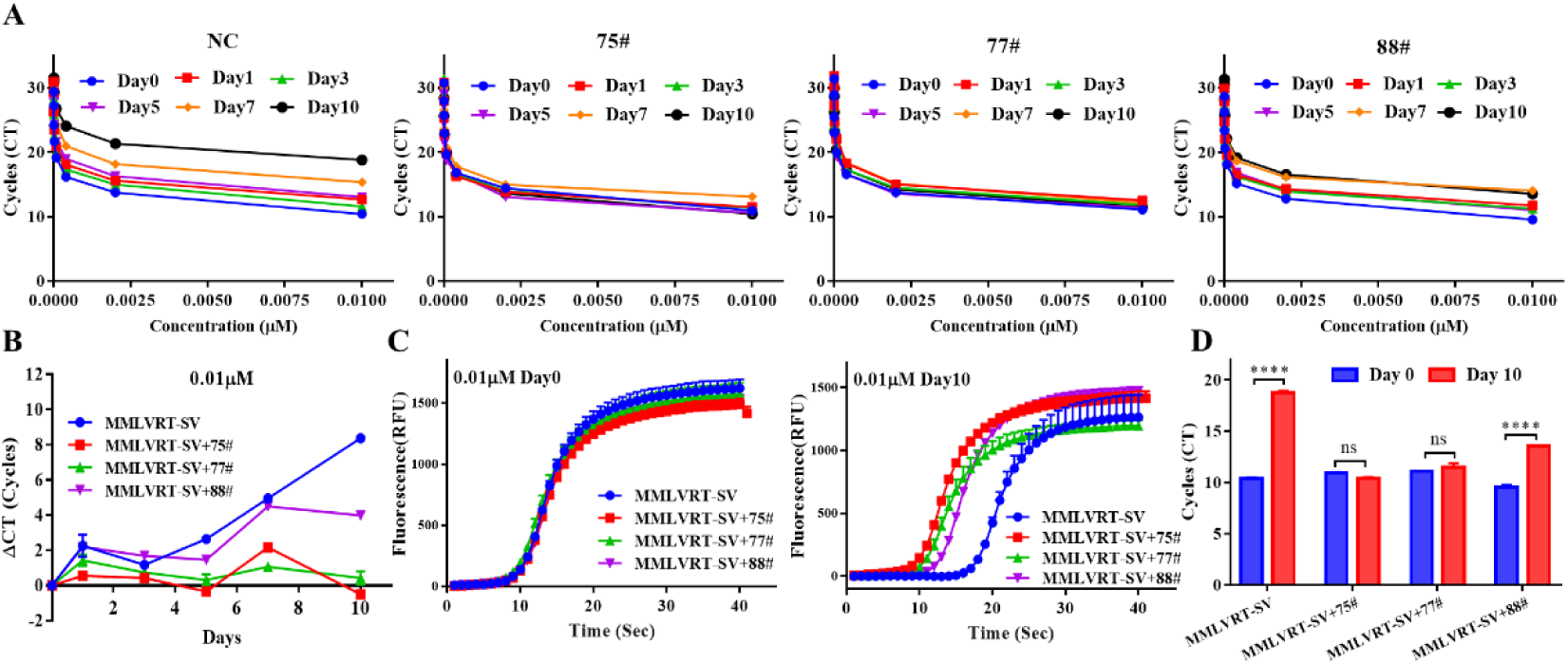
Binder75#, 77# and 88# enhance the long-term storage performance (efficiency and stability) of MMLV RT-SV. (A) RT-qPCR determination of the changes in the amplification cycle number of MMLV RT-SV at different storage time points with or without binder addition. (B) Under the condition of 0.01µM, the changes in the number of cycles of MMLV RT-SV in the RT-PCR process with or without binder addition as the number of storage days increases. (C) On day 0 and day 10, under the condition of 0.01µM, the fluorescence amplification curves of MMLV RT-SV in the RT-PCR process with or without binder addition. (D) On day 0 and day 10, under the condition of 0.01µM, the statistics of the cycle number of MMLV RT-SV in the RT - PCR process with or without binder addition.

### 2.6 The Binding Mode of Binder and the Core Sites in the Interaction Interface

To better verify the binding mode of the binder, we conducted an in-depth analysis of the interaction interface and key residues based on the complex structure predicted by AlphaFold3. The ipTM values of Binder75#, 77#, and 88# with MMLV RT-SV were 0.86, 0.80, and 0.84, respectively (Supplementary Figure 6). Structural analysis of the three candidate binders revealed that all binder bound to MMLV RT-SV according to the scaffold patterns we set. Binder 75# featured five α-helices, Binder 77# consisted of three nearly parallel α-helices, and Binder 88# formed a “V-shaped” structure with four α-helices (Supplementary Figure 7). All three binders exhibited strong negative surface charge (in terms of charge value and interaction area), which facilitated binding to the positively charged pocket of MMLV RT-SV (Figure 6A).

**Figure 6.**
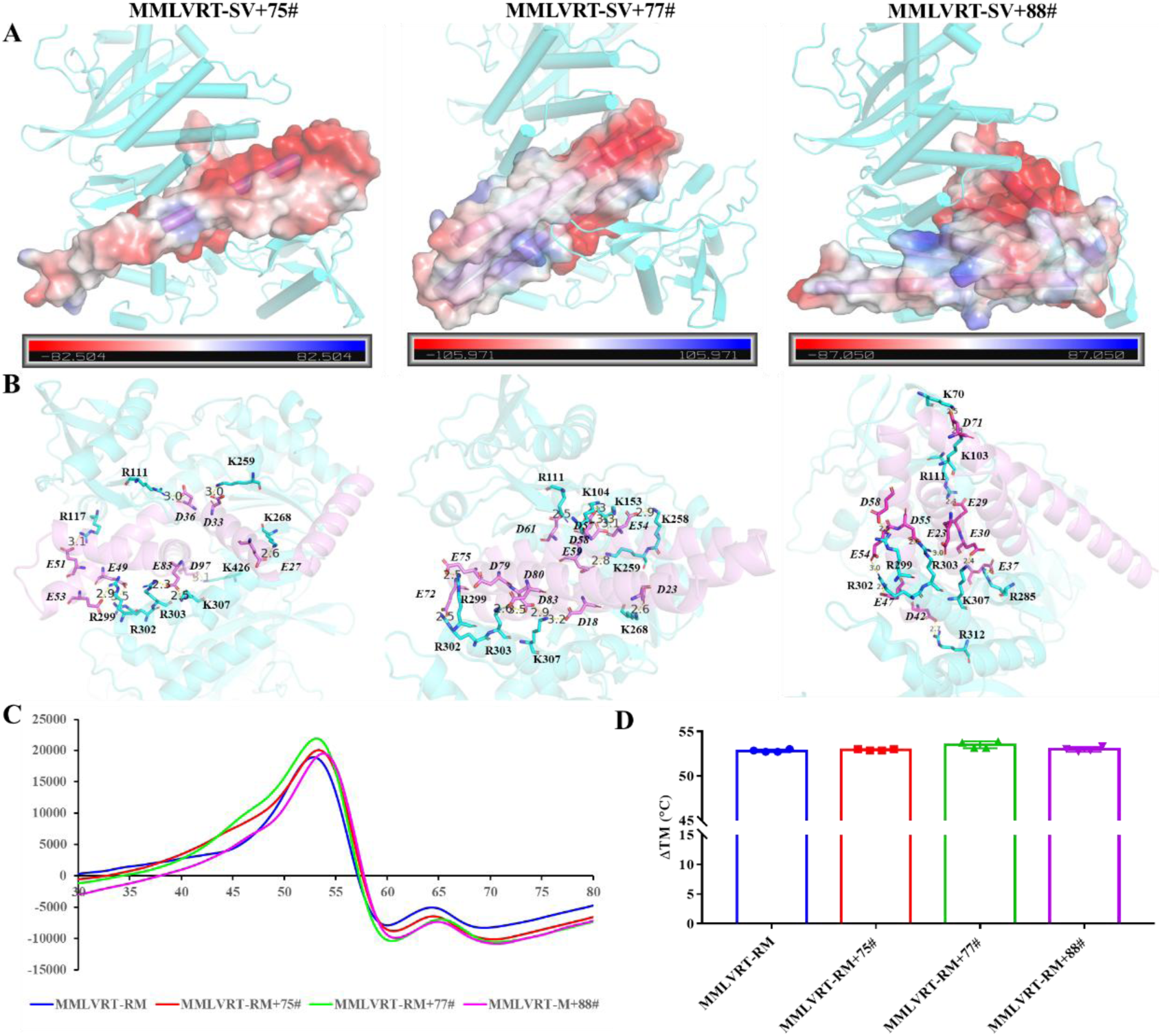
Interaction mode and analysis of the key sites of the complex structure of Binder75#, 77# and 88# with MMLV RT-SV. (A) AlphaFold3 predicted structures of the complexes of Binder75#, 77# and 88# with MMLV RT-SV and the display of the surface charge of the binders. (B) Analysis of the key residues in the interaction interface between Binder75#, 77# and 88# and MMLV RT-SV, with only the salt bridge interactions shown. The binder is in pink, and MMLV RT-SV is in light green. (C-D) DSF identification of the thermal denaturation curves and solubility temperature statistics of a mixture of 0.74 μM MMLV RT-RM and 4.4 μM Binder.

The interface areas of Binder75#, 77#, and 88# with MMLV RT-SV were 2314.4 Å², 2284.6 Å², and 2323.6 Å², respectively (Supplementary Table 3). The solvation free energy gain upon formation of the interface (Δ^i^G) was -17.2 kcal/mol, -5.9 kcal/mol, and -7.2 kcal/mol, respectively, with P-values all greater than 0.75. In the interaction interface, Binder75#, 77#, and 88# formed 32, 28, and 33 hydrogen bonds and 16, 25, and 35 salt bridges with MMLV RT-SV, respectively (Supplementary Tables 3). Further in-depth analysis of the interaction sites revealed that Binder75#, 77#, and 88# all shared three core interacting amino acid sites: R303, R111, and K307, all located in the nucleic acid-binding pocket of MMLV RT-SV (Figure 6B, Supplementary Tables 4-6). Therefore, we further constructed revertant mutants of MMLV RT-SV (MMLV RT-RM, R303A, R111A, and K307A) and used DSF experiments to investigate whether MMLV RT-RM could still bind to Binder75#, 77#, and 88#, to verify whether the binder bound to the nucleic acid-binding pocket of MMLV RT-SV as previously designed. The results showed that the melting temperature of MMLV RT- RM-binder was consistent with that of MMLV RT-RM, both at 53℃ (Figure 6C&D). This indicates that MMLV RT-RM lost its binding ability to the binder, and the three sites R303, R111, and K307 in the nucleic acid-binding pocket of MMLV RT-SV are the key sites for the binding of Binder75#, 77#, and 88# to MMLV RT-SV. These data indicate that Binder75#, 77#, and 88# bind to the nucleic acid-binding pocket centered on R303, R111, and K307, thereby enhancing the thermal stability and long-term storage performance of MMLV RT-SV.

## 3. Discussion

This study pioneers a synergistic approach combining multi-site mutagenesis and AI-driven de novo protein design to overcome the long-standing stability-activity trade-off in MMLV reverse transcriptase engineering. By targeting the positively charged nucleic acid-binding pocket, we engineered the high-stability variant MMLV RT-SV and rationally designed three negatively charged binders that stabilize the enzyme via electrostatic complementarity. This dual strategy not only enhances thermostability and storage resilience but also preserves catalytic function, offering a transformative solution for applications requiring robust enzymatic performance under demanding conditions.

The success of this approach lies in the precise mimicry of natural nucleic acid-enzyme interactions. Traditional stabilization strategies, such as rigidifying mutations or directed evolution, often inadvertently disrupt functional dynamics or substrate accessibility. In contrast, our de novo binders act as “electrostatic clamps,” reinforcing the structural integrity of the nucleic acid-binding pocket without occluding the catalytic core. This is evidenced by the competitive displacement of binders upon T/P addition, which ensures uninterrupted reverse transcription while maintaining stability benefits. The concentration-dependent Tm enhancement and nanomolar affinities further validate the design principle that targeted charge complementarity can achieve both high specificity and stabilization efficacy. Notably, the differential performance of the binders (e.g., Binder 88# showing highest affinity but moderate storage stability) suggests a balance between binding strength and functional plasticity that warrants further exploration.

This work challenges the conventional paradigm of enzyme stabilization through direct modification. While prior studies have utilized chaperones or chemical additives, our AI-designed binders represent a paradigm shift by providing genetically encodable, customizable stabilizers. The 8°C Tm increase in the MMLV RT-SV-binder complex surpasses most reported improvements from single-point mutations, with the added advantage of eliminating mutagenesis-induced activity loss. Furthermore, the retention of reverse transcription efficiency after 10-day storage at 37°C demonstrates unprecedented long-term stability—a critical advancement for diagnostic reagents and biomanufacturing workflows where cold-chain logistics are impractical.

The broader implications extend beyond reverse transcriptase optimization. We establish a generalizable framework for stabilizing nucleic acid-processing enzymes through charge-complementary binder design, applicable to DNA polymerases, CRISPR-associated proteins, and RNA-modifying enzymes. The identification of R303, R111, and K307 as critical interface residues provides a blueprint for future binder optimizations. However, limitations such as potential immunogenicity in therapeutic contexts or scalability of binder co-expression in industrial settings must be addressed. Future studies should explore fusion protein architectures and evaluate performance in complex biological matrices to fully realize the technology’s potential.

By merging computational biophysics with functional validation, this study redefines enzyme engineering possibilities. It highlights how AI-powered design can circumvent evolutionary constraints, offering a rapid, rational path to bespoke enzyme stabilizers. As molecular diagnostics and synthetic biology increasingly demand enzymes functioning under extreme conditions, our strategy provides a versatile toolkit to meet these challenges while preserving catalytic fidelity.

## 3. Materials and methods

### 3.1 Design and Optimization of MMLVRT-SV and Binders

The process for designing Binder for MMLVRT-SV is as follows:

1. Candidate Scaffold Generation and Preliminary Screening: RFdiffusion was used to generate over 200 potential MMLVRT-SV Binder scaffolds. PDBePISA was used to calculate MMLVRT-SV-Binder interaction areas, topological structures, etc., to screen scaffolds with binding potential.
2. Sequence Optimization and Restoration: After scaffold generation, sequences of Binder were optimized using ProteinMPNN. This method balanced sequence diversity, thermal stability, and functionality.
3. Molecular Conformation Exploration and Binding Pattern Optimization: AlphaFold3 model was used to comprehensively explore molecular conformations of candidate Binder with MMLVRT-SV.
4. Folding Consistency and Structural Validation: Using AlphaFold3, the folding consistency of Binder monomers and their complexes was assessed to ensure stable and reliable binding patterns.
5. Stability and Expression Probability Assessment: Energy optimization and scoring functions of Rosetta and MPNNsol model were used to systematically evaluate the structural stability and expression probability of candidate Binder.

### 3.2 Expression and Purification of MMLVRT-SV

The full-length gene-synthesized heat-resistant MMLV reverse transcriptase mutants were transformed into *E. coli* BL21 Star (DE3) cells. After plate culture, a single colony was picked and inoculated into LB liquid medium containing 100 μg/mL ampicillin. When the OD₆₀₀ of the culture reached 0.6-0.8, 1.0 mmol/L IPTG was added to induce expression at 25°C for 5 hours. The cells were then harvested by centrifugation at 4°C, 4200 rpm, and resuspended in lysis buffer (50 mM Tris-HCl, 300 mM NaCl, 20 mM imidazole, 5% glycerol, pH 7.8) followed by sonication. After centrifugation at 4°C, 12000 rpm for 30 minutes to remove cell debris, the supernatant was loaded onto a pre-equilibrated Ni-NTA affinity chromatography column (equilibration buffer components were the same as lysis buffer). The target protein was eluted with a gradient elution buffer containing 250 mM imidazole (other components were the same as equilibration buffer). After SDS-PAGE analysis, the target fractions were pooled and dialyzed in two steps with 50 mM Tris-HCl buffer (pH 8.0): first dialysis for 6 hours, followed by overnight dialysis (both at 4°C).

The dialyzed product was further purified using an SP cation exchange column (equilibration buffer: 20 mM Tris-HCl, 200 mM NaCl, 5% glycerol, 1 mM DTT, 1 mM EDTA, pH 7.8) and eluted with a gradient elution buffer containing 600 mM NaCl. The purified fractions were collected and verified by SDS-PAGE, and the final product was stored in 50% glycerol. Protein purity was analyzed by 10% polyacrylamide gel SDS-PAGE, and protein concentration was quantified using the Coomassie Brilliant Blue colorimetric method.

### 3.3 Expression and Purification of Binders

The full-length gene-synthesized binders expression vector pET32a- 6His-sumo-Binders was transformed into *E. coli* BL21 Star (DE3) cells. After activation by streaking on plates, a single colony was picked and inoculated into LB liquid medium containing 100 μg/mL ampicillin for expansion culture. When the OD₆₀₀ of the culture reached 0.6-0.8, 0.4 mmol/L IPTG was added to induce protein expression at 25°C for 6 hours. The bacterial culture was then harvested by centrifugation at 4°C, 4200 rpm for 10 minutes. The cells were resuspended in pre-cooled lysis buffer (50 mM Tris-HCl, 300 mM NaCl, 20 mM imidazole, 5% glycerol, pH 7.8) and sonicated on ice (parameters: 200 W, 3 seconds sonication/5 seconds interval, total duration 30 minutes). After centrifugation to collect the supernatant containing the crude protein extract, the crude extract was loaded onto a Ni-NTA metal chelating chromatography column pre-equilibrated with equilibration buffer (same as lysis buffer). The target protein was eluted with a linear gradient elution buffer containing 250 mM imidazole (other components were the same as equilibration buffer). After collecting the target protein peak, ULP1 protease was added at a ratio of 1:10 (ULP1: protein solution) for SUMO tag cleavage, and the reaction was carried out overnight (16 hours) at 4°C. The cleavage products were further purified using a Q anion exchange column (equilibration buffer: 20 mM Tris-HCl, 200 mM NaCl, 5% glycerol, 1 mM DTT, 1 mM EDTA, pH 7.8) and eluted with a gradient elution buffer containing 600 mM NaCl. The highly pure fractions verified by SDS-PAGE were collected and stored in 50% glycerol. Protein purity was analyzed by 16.8% tricine-SDS-PAGE, and protein concentration was quantified using the Coomassie Brilliant Blue staining method and standard curve.

### 3.3 Differential Scanning Fluorimetry (DSF) for Assessing Protein Thermal Stability

Differential Scanning Fluorimetry (DSF) was used to assess protein thermal stability by slowly heating samples in a real-time PCR instrument and measuring the binding of fluorescent dye to structurally altered proteins. The procedure was as follows: First, protein samples were diluted to 250-600 ng/μL with reaction buffer (250 mM Tris-HCl pH 8.3, 375 mM KCl, 15 mM MgCl₂, 50 mM DTT). Then for the T/P test, Primer (5’- TGGAATCAGGTGTCGCACTCTG-3’) and the template (5’- rArArCrArGrArGrUrGrCrGrArCrArCrCrUrGrArUrUrCrCrArU- 3’) were each diluted to 100 μM, mixed in a 1:1 ratio, 60℃, 5min, and annealed to form a complex. The T/P complex is then mixed 1:1 with the protein (MMLV or MMLV-BINDER) to produce a protein DSF sample. For the DSF reaction, a 20 μL mixture was prepared containing 16 μL DSF buffer (20 mM HEPES, 150 mM NaCl, pH 7.4), 2 μL protein sample, and 2 μL 20 × AGSYPRO Orange dye(AG31002)(DMSO diluted; purchased from the ACCURATE BIOTECHNOLOGY(HUNAN)CO.,LTD, ChangSha, China). The reaction program was set as follows: pre-incubation at 8°C for 2 minutes, pre-reaction at 25°C for 2 minutes, followed by heating from 25°C to 99°C at a rate of 0.05°C/10 seconds, with fluorescence signals collected via the X1M3 channel using a QuantStudio real-time PCR instrument. Finally, data were analyzed using the instrument’s software (protein thermal shift software) to calculate the melting temperature (Tm) and related parameters of the protein samples.

### 3.4 Biolayer interferometry (BLI) measurements

Purified Target MMLVRT-SV proteins were biotinylated using a biotinylation kit (21343, Thermo Fisher Scientific). Briefly, MMLVRT-SV and biotinylation reagent were mixed with 1:1 molar ratio in PBS at room temperature. The reaction mixtures were incubated at room temperature for 1 h, which was then dialyzed using Thermo Scientific Pierce Zeba (89882, Thermo Fisher Scientific) to remove unreacted biotinylation reagent.

BLI experiments were performed using a GatorPrime instrument from GatorBio. All assays were run at 30 ℃ with continuous 1000 rpm shaking. The assay buffer consisted of PBS, 0.02% Tween-20, adjusted to pH 7.5. Biotinylated MMLVRT-SV proteins were tethered on streptavidin (SA-XT) biosensors (GatorBio) by dipping sensors into protein solutions. SA-XT biosensors were then washed in assay buffer for 120 s to eliminate nonspecifically bound protein and establish stable baselines. Then SA-XT biosensor dipped into various concentrations of Binder75#(0.375 to 6 μM), Binder77#(0.312 to 5 μM)and Binder88# (31.25-500 nM) for 200 s to record the association phase. Then, the sensors were placed into wells containing measuring buffer for 200 s to record the dissociation phase. All the data were analyzed by Gator Bio data analysis software. The equilibrium dissociation constant (Kd) values were calculated from the ratio of Koff to Kon based on global fitting (1:1 binding model) of several curves generated from serial dilutions of the proteins.

### 3.5 Performance Verification of MMLVRT-WT and MMLVRT-SV

optimized using 293T RNA as the template. Both MMLVRT-WT and MMLVRT-SV (250 ng each) were added to the reaction system, with reverse transcription temperatures increased to 62°C and 65°C. The resulting cDNA was amplified using (ACCURATE BIOTECHNOLOGY’s Exp Taq DNA Polymerase Ver.2 (AG11403), which contains high concentrations of Mg²⁺ and dNTPs. For the 4 kb and 10 kb target fragments, primer pairs P0377/P0378 and P0383/P0384 were used respectively, with extension times set at 4 minutes per cycle (4 kb) and 10 minutes per cycle (10 kb). The PCR program included 94℃ pre-denaturation for 2 minutes, 35 cycles (94℃ for 30 seconds, 55-60℃ annealing for 30 seconds, extension at 72℃ for 4 minutes), and a final extension at 72℃ for 10 minutes.

### 3.6 Inhibition of Reverse Transcription by MMLVRT-SV and Binder

Mix MMLVRT-SV and binder at a molar ratio of 1:6, and prepare a reaction system without Oligo dT (18T) Primer and dNTP Mix according to the ACCURATE BIOTECHNOLOGY’s Evo M-MLV RT Kit (AG11603). Incubate at 37°C for 30 or 10 minutes. The concentration gradient of ligo dT (18T) Primer are 0.125, 0.1, 0.075, 0.05, 0.025, 0.005 μM, and the concentration gradient of 293T RNA are 500, 100 ng. Amplify the cDNA using Exp Taq DNA Polymerase Ver.2 (AG11403) from the ACCURATE BIOTECHNOLOGY(HUNAN)CO., LTD, ChangSha, China), with primers P0377 and P0378 for the 4 kb target fragment.

### 3.7 RT-qPCR Validation Stability of Binder on MMLVRT-SV

The molar ratio of MMLV reverse transcriptase mutants and wild-type to binder was 1:6. After mixing, the mixture was incubated at 37℃ for 30 minutes. The incubated mixture was diluted to a working concentration of 90 ng/μL using MMLV storage buffer (containing 50% glycerol) and then store .

After storage, the enzyme samples were systematically diluted to achieve final concentrations of 0.01, 0.002, 0.0004, 0.00008, 0.000016, 0.0000032, 0.00000064, and 0.000000128 μmol/L. Reverse transcription was conducted using MS2 RNA as the template. Subsequently, quantitative PCR analysis was performed on the synthesized cDNA templates utilizing primers P0330 and P0329, with detection accomplished through the SYBR Green Pro Taq HS Pre-Mixed qPCR Kit AG11701 (Accurate Biotechnology). Amplification efficiency was evaluated by analyzing fluorescence intensity curves during thermal cycling, while cycle threshold (CT) values were determined for quantitative expression profiling.

## 4. Declarations

### Author Contributions

Tao Li and Yibo Zhu conceived and designed the study, sorted out all the experimental results, wrote the manuscript and responsible for the delivery and follow-up work of the article. Hongwei Liu, Fang Qu mainly completed the experimental content of this study. Hong Yao, Chunlei Ge Yingzhi Wang, JuanJuan Yao, Shumin You and Lin Hua assisted in the completion of structural analysis and other studies.

## Acknowledgements

Not applicable.

## Consent for publication

All authors have reviewed the final version of the manuscript and approved it for publication.

## Availability of data and materials

All data generated or analyzed during this study are included in this article and additional data are available from the corresponding author upon reasonable request.

## Supplementary Material

**Supplementary Figure 1.**
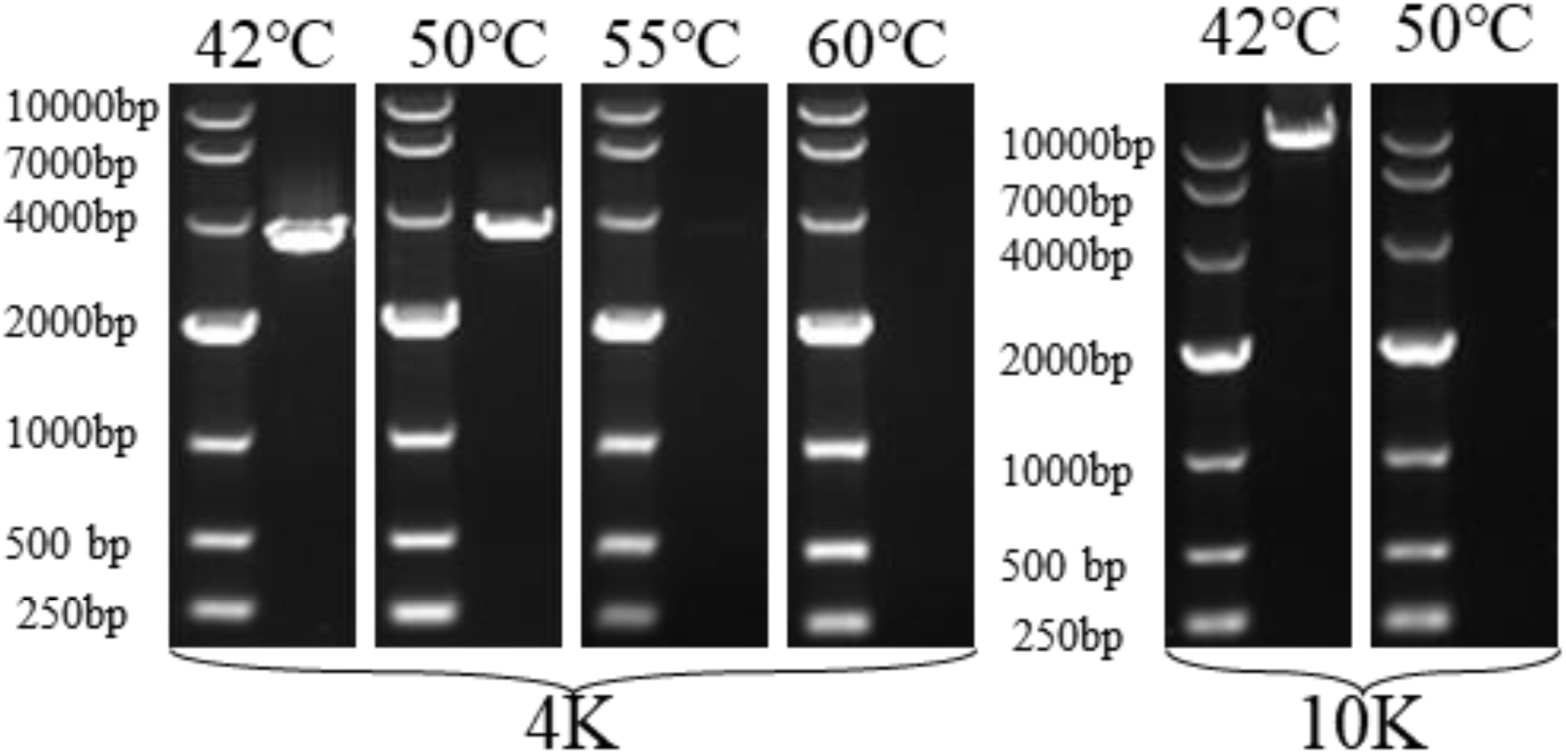
Agarose gel identification of reverse transcription results of MMLVRT-WT and MMLVRT-SV at different temperatures (42℃, 50℃, 55℃, 60℃) and with different molecular weight templates (4K and 10K).

**Supplementary Figure 2.**
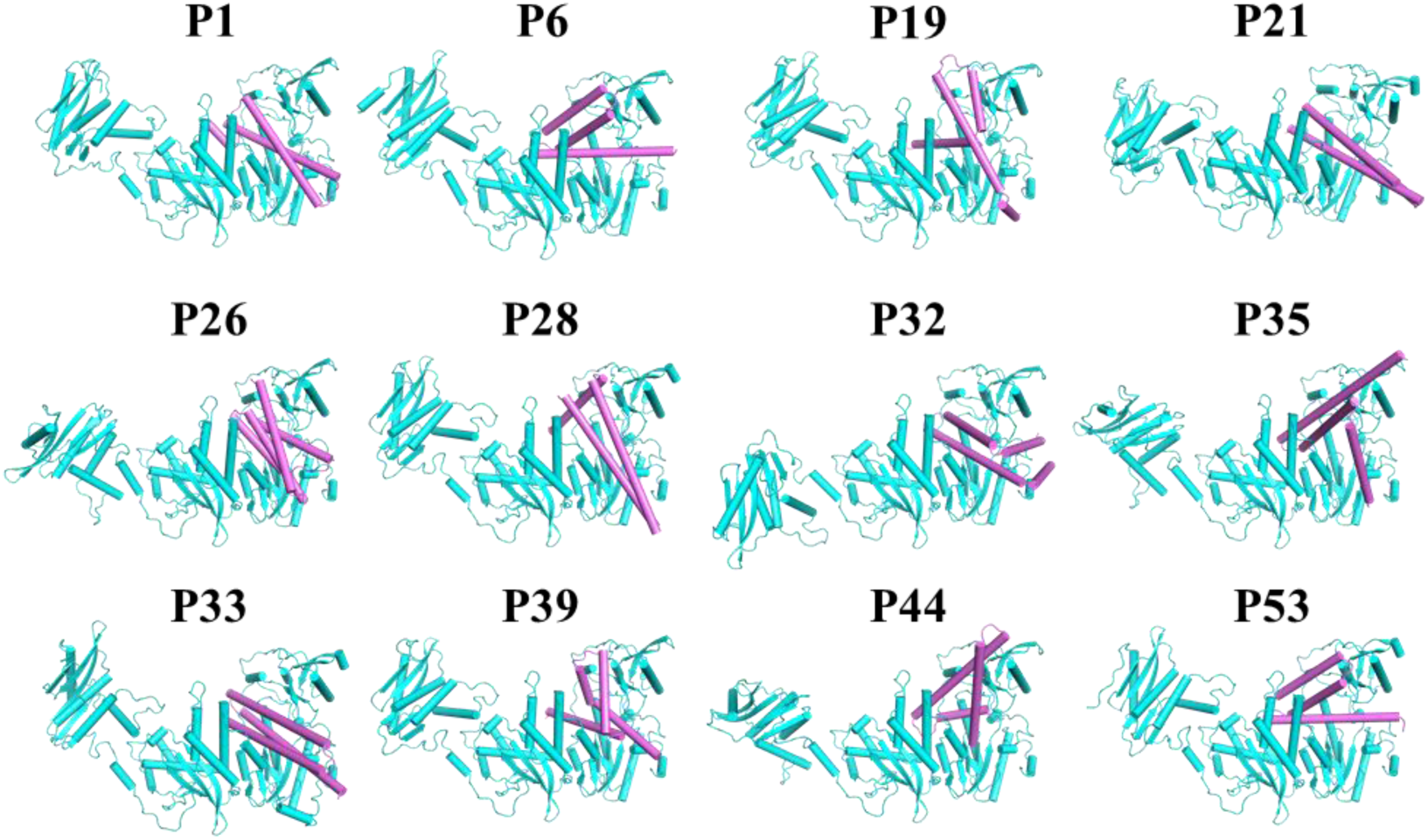
RFdiffusion platform for geometrically constrained pocket topology matching design. 12 high-matching scaffolds generated by diffusion models are shown in cartoon representation. MMLV RT-SV is colored blue, while the different scaffolds are colored pink. The sequences of these scaffolds contain 88 and 100 amino acids respectively.

**Supplementary Figure 3.**
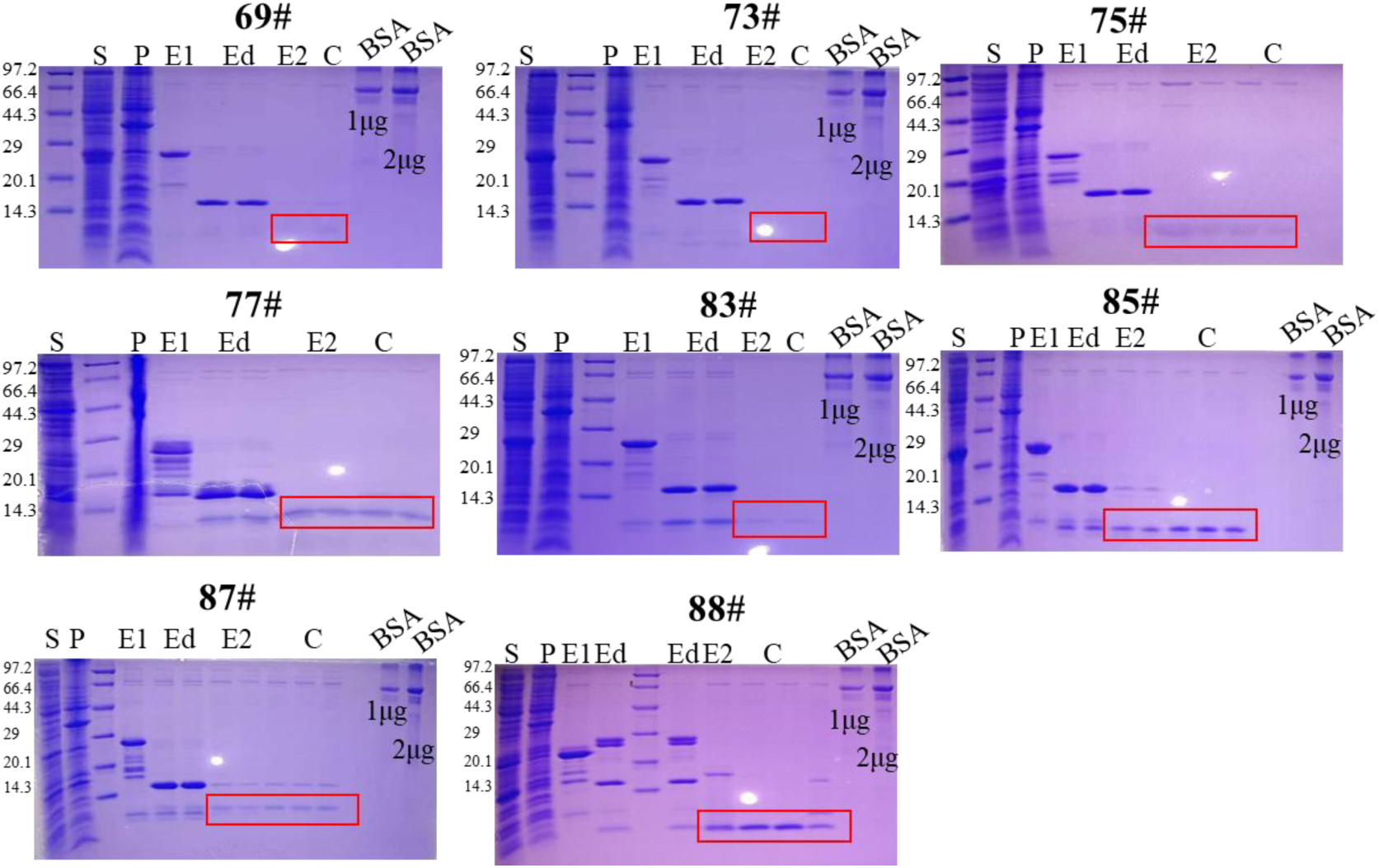
Figure. PAGE gel identification of binder expression and purification results. E: Supernatant, P: Precipitation, Ed: Enzymatic digestion, E1: First elution, E2: Second elution, C: Concentration, BSA: Bovine serum albumin. Note: The designed binder targets the strongly positively charged nucleic acid-binding pocket and displays a strong negative charge, resulting in a forward-shifted Coomassie Brilliant Blue-stained band. The red box indicates the binder position after SUMO-6His tag removal via enzymatic digestion.

**Supplementary Figure 4.**
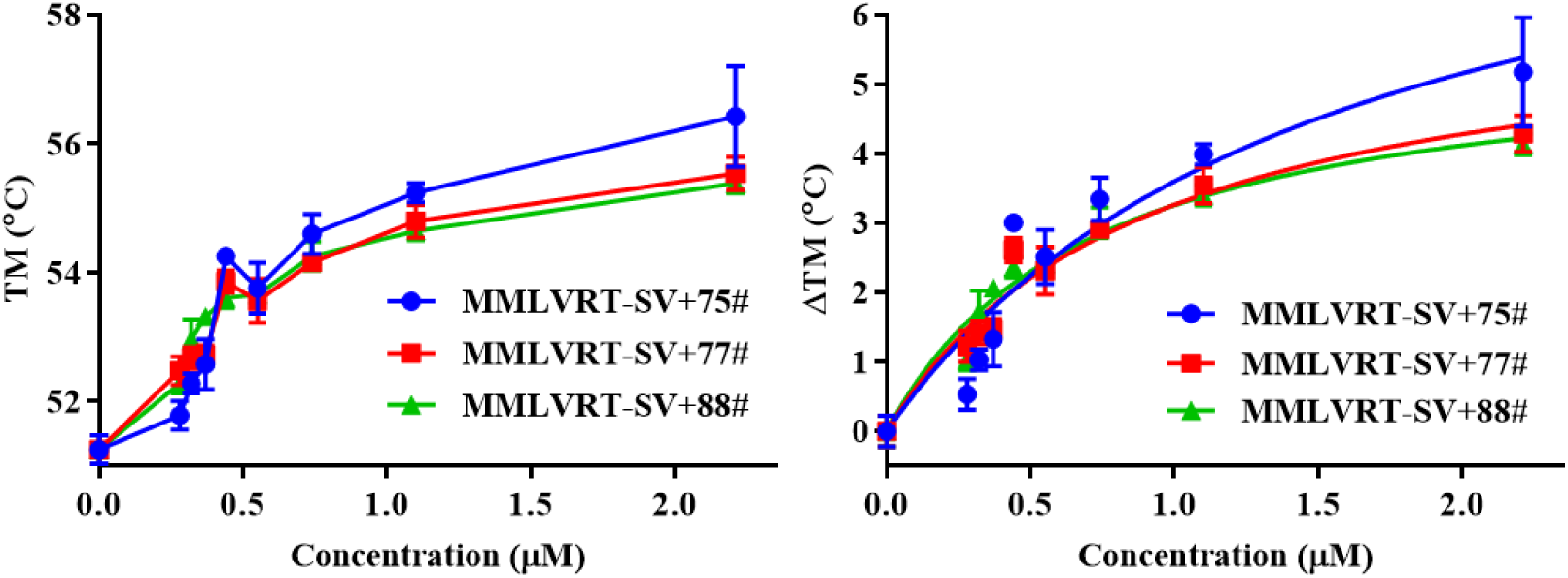
Statistical results of DSF determined changes in the thermal denaturation curves of MMLV RT-SV under gradient concentrations (0 to 2.21 μM) of Binder added.

**Supplementary Figure 5.**
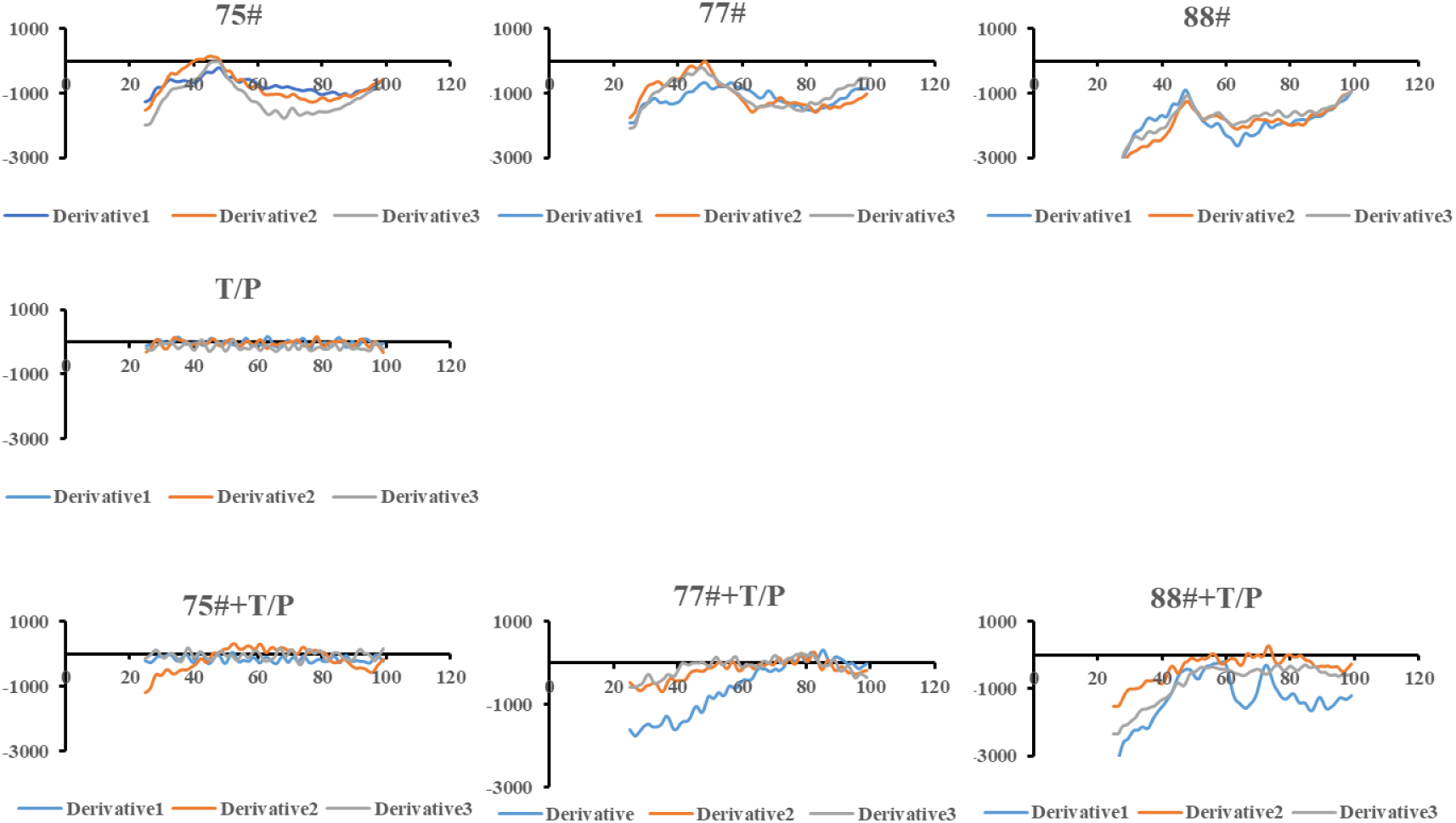
DSF experiments of the binder alone, T/P alone, and the Binder-T/P mixture. No significant signal values were detected, and the melting temperatures could not be evaluated.

**Supplementary Figure 6.**
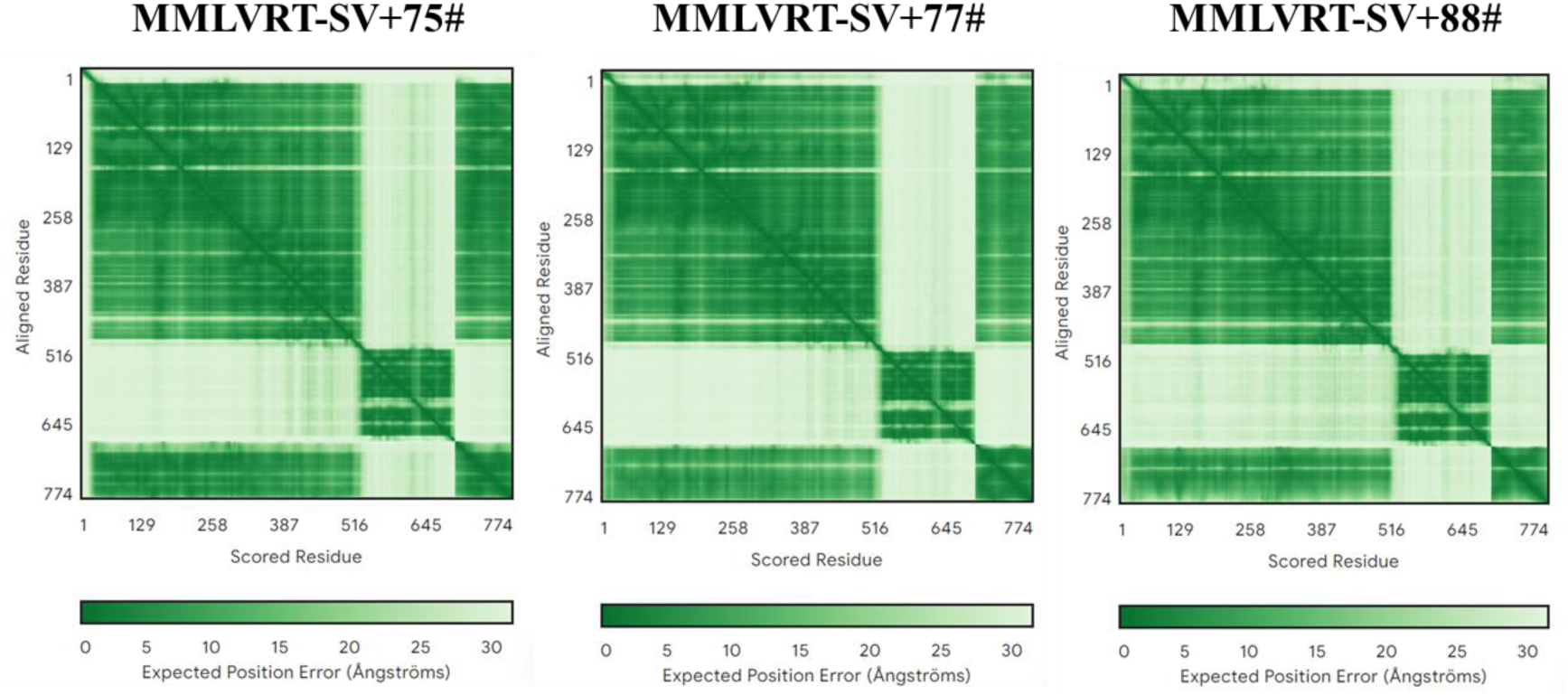
AlphaFold3-predicted structure of the protein with confidence score (pLDDT) color-coded in green. The green color indicates regions with higher prediction confidence.

**Supplementary Figure 7.**
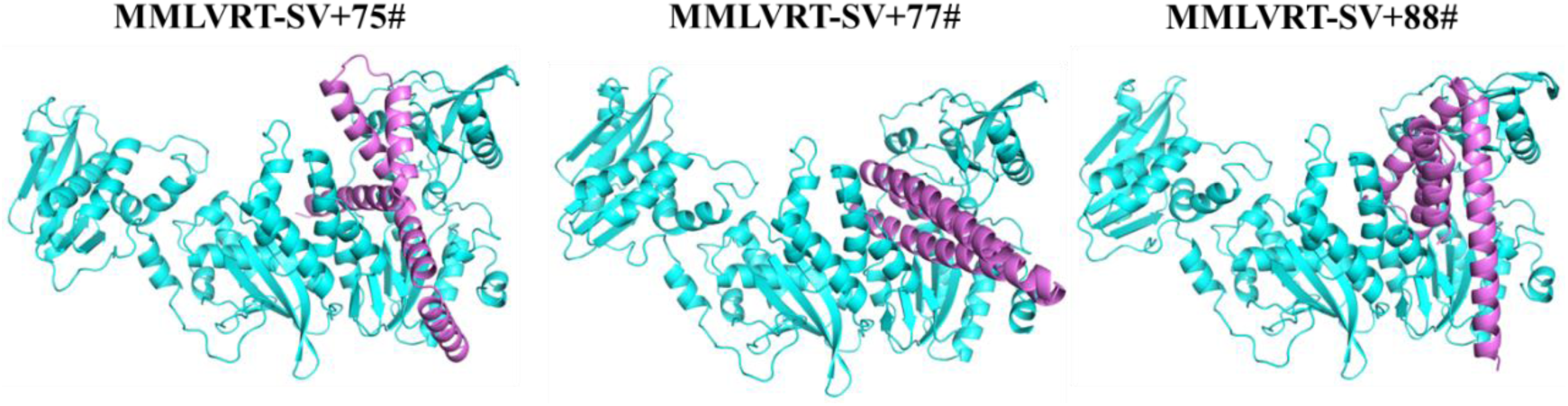
Predicted complex structures of Binder75#, 77#, and 88# with MMLV RT-SV by AlphaFold3. MMLV RT-SV is colored blue, and the binders are colored pink.

**Supplementary Table 1.**
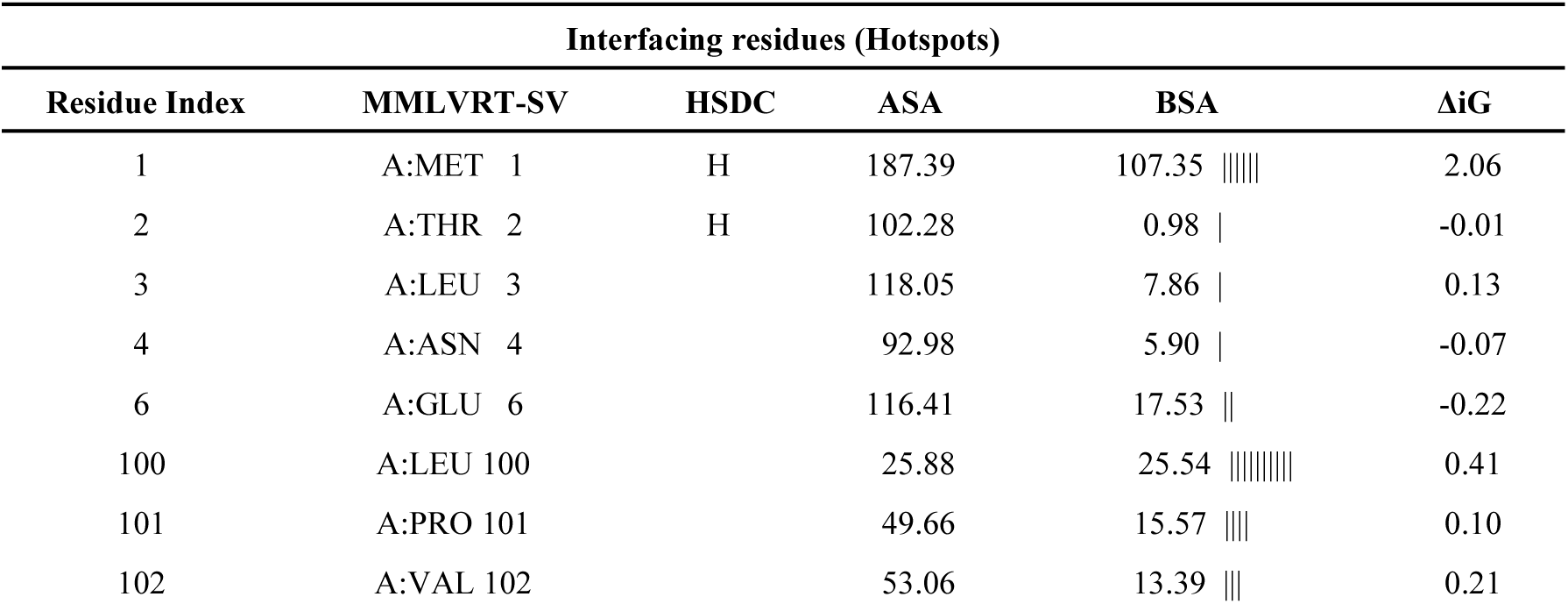
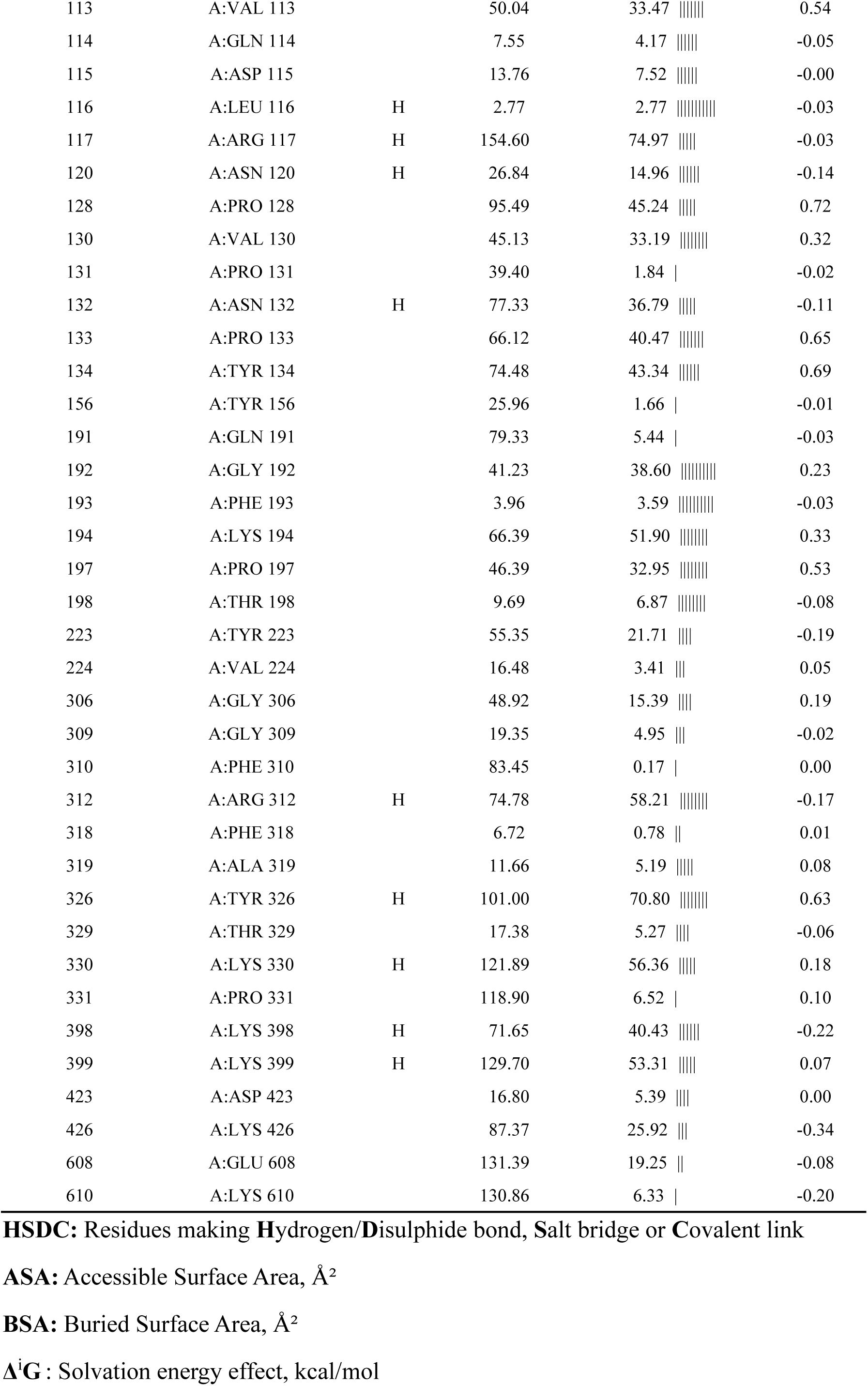
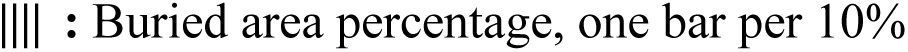
Key interaction sites between MMLVRT-SV and T/P through PDBePISA.

**Supplementary Table 2.** Physical parameters and expression statistics of Binders.

| Name | MW<br>(Kda) | ABS | Bacterial<br>Biomass(g) | OD280<br>(mg/ml) | OD280/abs<br>(ng/ul) | Volume<br>(ml) |
| --- | --- | --- | --- | --- | --- | --- |
| ligand69+sumo | 10 | 0.297 | 3 | 0.47 | 1582.49 | 10.5 |
| ligand73+sumo | 10 | 0.293 | 2.5 | 0.37 | 1262.80 | 10 |
| ligand75+sumo | 10 | 0.3 | 2.57 | 0.34 | 1133.33 | 7.5 |
| ligand77+sumo | 10.5 | 1.48 | 2.49 | 1.03 | 695.95 | 6 |
| Ligand83+sumo | 10 | 0.568 | 2.5 | 0.42 | 739.44 | 7.5 |
| Ligand85+sumo | 8.3 | 1.018 | 3.27 | 2.06 | 2023.58 | 2.5 |
| Ligand87+sumo | 8.4 | 1.821 | 2.99 | 2.06 | 1131.25 | 2.5 |
| ligand88+sumo | 10.7 | 1.31 | 2.73 | 1.47 | 1122.14 | 7.5 |

**Supplementary Table 3.** Interaction Parameters between Binders and MMLVRT-SV.

| Interface Interaction Parameter |  |  |  |  |  |  |  |  |  |  |  |  |  |  |
| --- | --- | --- | --- | --- | --- | --- | --- | --- | --- | --- | --- | --- | --- | --- |
| ## | Structure 1 | | | | Structure 2 | | | | Interface<br>area, Å<sup>2</sup> | $\Delta^i G$<br>kcal/mol | $\Delta^i G$<br>P-value | N<sub>HB</sub> | N<sub>SB</sub> | N<sub>DS</sub> |
| NN | Range | <sup>i</sup>N<sub>at</sub> | <sup>i</sup>N<sub>res</sub> | Surface<br>Å<sup>2</sup> | Range | <sup>i</sup>N<sub>at</sub> | <sup>i</sup>N<sub>res</sub> | Surface<br>Å<sup>2</sup> |  |  |  |  |  |  |
| 75# | A | 252 | 73 | 35114 | B | 214 | 59 | 7131 | 2314.4 | -17.2 | 0.765 | 32 | 17 | 0 |
| 77# | A | 243 | 64 | 35184 | B | 226 | 54 | 6311 | 2284.6 | -5.9 | 0.835 | 28 | 25 | 0 |
| 88# | A | 242 | 67 | 32756 | B | 216 | 51 | 7098 | 2323.6 | -7.2 | 0.822 | 33 | 35 | 0 |

**Supplementary Table 4.**
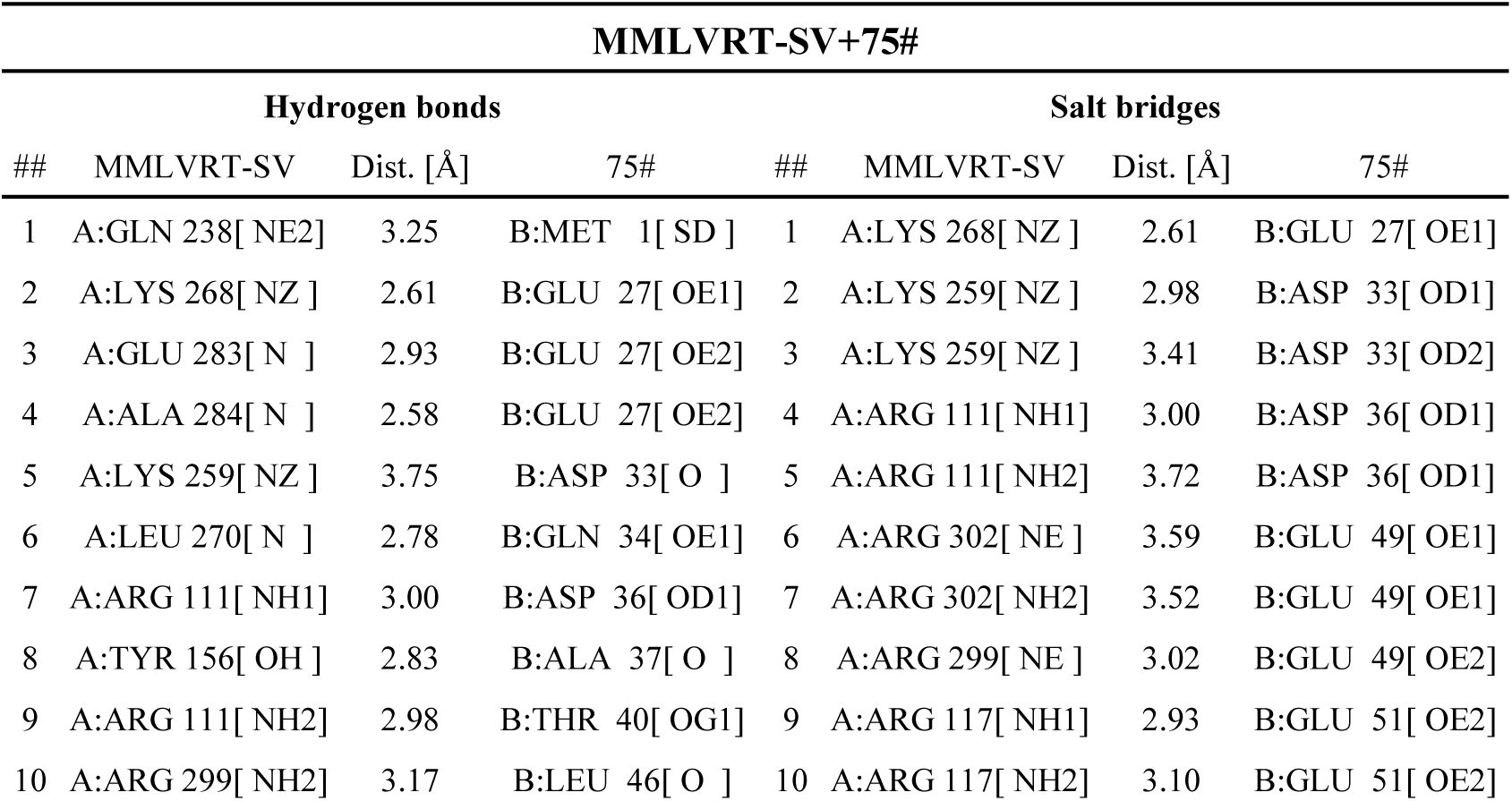
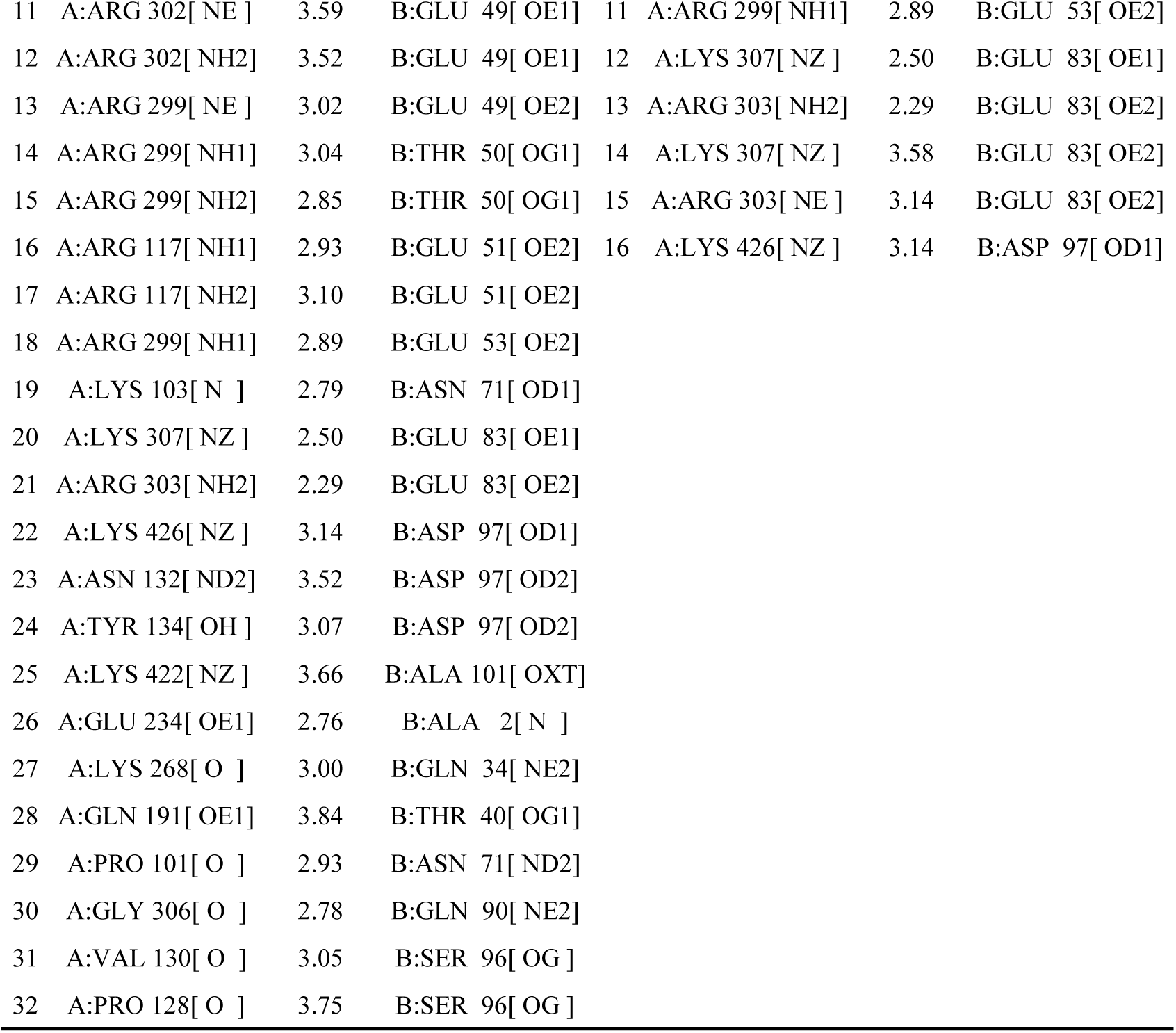
Key Residues and Interaction Statistics between Binder75# and MMLVRT-SV.

| MMLVRT-SV+75# |  |  |  |  |  |  |  |  |
| --- | --- | --- | --- | --- | --- | --- | --- | --- |
| Hydrogen bonds |  |  |  |  | Salt bridges |  |  |  |
| ## | MMLVRT-SV | Dist. [Å] | 75# |  | ## | MMLVRT-SV | Dist. [Å] | 75# |
| 1 | A:GLN 238[ NE2 ] | 3.25 | B:MET 1[ SD ] |  | 1 | A:LYS 268[ NZ ] | 2.61 | B:GLU 27[ OE1 ] |
| 2 | A:LYS 268[ NZ ] | 2.61 | B:GLU 27[ OE1 ] |  | 2 | A:LYS 259[ NZ ] | 2.98 | B:ASP 33[ OD1 ] |
| 3 | A:GLU 283[ N ] | 2.93 | B:GLU 27[ OE2 ] |  | 3 | A:LYS 259[ NZ ] | 3.41 | B:ASP 33[ OD2 ] |
| 4 | A:ALA 284[ N ] | 2.58 | B:GLU 27[ OE2 ] |  | 4 | A:ARG 111[ NH1 ] | 3.00 | B:ASP 36[ OD1 ] |
| 5 | A:LYS 259[ NZ ] | 3.75 | B:ASP 33[ O ] |  | 5 | A:ARG 111[ NH2 ] | 3.72 | B:ASP 36[ OD1 ] |
| 6 | A:LEU 270[ N ] | 2.78 | B:GLN 34[ OE1 ] |  | 6 | A:ARG 302[ NE ] | 3.59 | B:GLU 49[ OE1 ] |
| 7 | A:ARG 111[ NH1 ] | 3.00 | B:ASP 36[ OD1 ] |  | 7 | A:ARG 302[ NH2 ] | 3.52 | B:GLU 49[ OE1 ] |
| 8 | A:TYR 156[ OH ] | 2.83 | B:ALA 37[ O ] |  | 8 | A:ARG 299[ NE ] | 3.02 | B:GLU 49[ OE2 ] |
| 9 | A:ARG 111[ NH2 ] | 2.98 | B:THR 40[ OG1 ] |  | 9 | A:ARG 117[ NH1 ] | 2.93 | B:GLU 51[ OE2 ] |
| 10 | A:ARG 299[ NH2 ] | 3.17 | B:LEU 46[ O ] |  | 10 | A:ARG 117[ NH2 ] | 3.10 | B:GLU 51[ OE2 ] |

|  |  |  |  |  |  |  |  |
| --- | --- | --- | --- | --- | --- | --- | --- |
| 11 | A:ARG 302[ NE ] | 3.59 | B:GLU 49[ OE1] | 11 | A:ARG 299[ NH1] | 2.89 | B:GLU 53[ OE2] |
| 12 | A:ARG 302[ NH2] | 3.52 | B:GLU 49[ OE1] | 12 | A:LYS 307[ NZ ] | 2.50 | B:GLU 83[ OE1] |
| 13 | A:ARG 299[ NE ] | 3.02 | B:GLU 49[ OE2] | 13 | A:ARG 303[ NH2] | 2.29 | B:GLU 83[ OE2] |
| 14 | A:ARG 299[ NH1] | 3.04 | B:THR 50[ OG1] | 14 | A:LYS 307[ NZ ] | 3.58 | B:GLU 83[ OE2] |
| 15 | A:ARG 299[ NH2] | 2.85 | B:THR 50[ OG1] | 15 | A:ARG 303[ NE ] | 3.14 | B:GLU 83[ OE2] |
| 16 | A:ARG 117[ NH1] | 2.93 | B:GLU 51[ OE2] | 16 | A:LYS 426[ NZ ] | 3.14 | B:ASP 97[ OD1] |
| 17 | A:ARG 117[ NH2] | 3.10 | B:GLU 51[ OE2] |  |  |  |  |
| 18 | A:ARG 299[ NH1] | 2.89 | B:GLU 53[ OE2] |  |  |  |  |
| 19 | A:LYS 103[ N ] | 2.79 | B:ASN 71[ OD1] |  |  |  |  |
| 20 | A:LYS 307[ NZ ] | 2.50 | B:GLU 83[ OE1] |  |  |  |  |
| 21 | A:ARG 303[ NH2] | 2.29 | B:GLU 83[ OE2] |  |  |  |  |
| 22 | A:LYS 426[ NZ ] | 3.14 | B:ASP 97[ OD1] |  |  |  |  |
| 23 | A:ASN 132[ ND2] | 3.52 | B:ASP 97[ OD2] |  |  |  |  |
| 24 | A:TYR 134[ OH ] | 3.07 | B:ASP 97[ OD2] |  |  |  |  |
| 25 | A:LYS 422[ NZ ] | 3.66 | B:ALA 101[ OXT] |  |  |  |  |
| 26 | A:GLU 234[ OE1] | 2.76 | B:ALA 2[ N ] |  |  |  |  |
| 27 | A:LYS 268[ O ] | 3.00 | B:GLN 34[ NE2] |  |  |  |  |
| 28 | A:GLN 191[ OE1] | 3.84 | B:THR 40[ OG1] |  |  |  |  |
| 29 | A:PRO 101[ O ] | 2.93 | B:ASN 71[ ND2] |  |  |  |  |
| 30 | A:GLY 306[ O ] | 2.78 | B:GLN 90[ NE2] |  |  |  |  |
| 31 | A:VAL 130[ O ] | 3.05 | B:SER 96[ OG ] |  |  |  |  |
| 32 | A:PRO 128[ O ] | 3.75 | B:SER 96[ OG ] |  |  |  |  |

**Supplementary Table 5.** Key Residues and Interaction Statistics between Binder77# and MMLVRT-SV.

| MMLVRT-SV +77# |  |  |  |  |  |  |  |
| --- | --- | --- | --- | --- | --- | --- | --- |
| Hydrogen bonds |  |  |  | Salt bridges |  |  |  |
| ## | MMLVRT-SV | Dist. [Å] | 77# | ## | MMLVRT-SV | Dist. [Å] | 77# |
| 1 | A:GLY 271[ N ] | 2.98 | B:GLU 16[ OE1] | 1 | A:LYS 307[ NZ ] | 3.70 | B:ASP 18[ OD1] |
| 2 | A:LYS 307[ NZ ] | 3.16 | B:ASP 18[ OD2] | 2 | A:LYS 307[ NZ ] | 3.16 | B:ASP 18[ OD2] |
| 3 | A:THR 282[ OG1] | 2.75 | B:ASP 19[ OD1] | 3 | A:LYS 268[ NZ ] | 2.64 | B:GLU 23[ OE1] |
| 4 | A:ARG 285[ N ] | 3.90 | B:ASP 19[ OD1] | 4 | A:LYS 258[ NZ ] | 2.89 | B:ASP 54[ OD2] |
| 5 | A:LYS 268[ NZ ] | 2.64 | B:GLU 23[ OE1] | 5 | A:LYS 104[ NZ ] | 3.52 | B:ASP 57[ OD1] |
| 6 | A:LYS 258[ NZ ] | 2.89 | B:ASP 54[ OD2] | 6 | A:ARG 111[ NH1] | 3.30 | B:ASP 57[ OD1] |
| 7 | A:ARG 111[ NH1] | 2.49 | B:ASP 57[ O ] | 7 | A:LYS 104[ NZ ] | 2.28 | B:ASP 57[ OD2] |
| 8 | A:ARG 111[ NH1] | 3.30 | B:ASP 57[ OD1] | 8 | A:LYS 153[ NZ ] | 3.26 | B:ASP 58[ OD1] |
| 9 | A:LYS 104[ NZ ] | 2.28 | B:ASP 57[ OD2] | 9 | A:LYS 259[ NZ ] | 3.71 | B:ASP 58[ OD2] |
| 10 | A:LYS 153[ NZ ] | 3.26 | B:ASP 58[ OD1] | 10 | A:LYS 153[ NZ ] | 3.10 | B:ASP 58[ OD2] |
| 11 | A:LYS 259[ NZ ] | 3.71 | B:ASP 58[ OD2] | 11 | A:LYS 259[ NZ ] | 2.80 | B:GLU 59[ OE1] |
| 12 | A:LYS 153[ NZ ] | 3.10 | B:ASP 58[ OD2] | 12 | A:ARG 111[ NH2] | 2.61 | B:ASP 61[ OD1] |
| 13 | A:LYS 259[ NZ ] | 2.80 | B:GLU 59[ OE1] | 13 | A:ARG 111[ NH2] | 2.49 | B:ASP 61[ OD2] |
| 14 | A:ARG 111[ NH2 | 2.49 | B:ASP 61[ OD2] | 14 | A:ARG 111[ NH1] | 3.76 | B:ASP 61[ OD2] |
| 15 | A:ASN 120[ ND2] | 3.82 | B:ALA 68[ O ] | 15 | A:ARG 302[ NH1] | 2.47 | B:GLU 72[ OE1] |
| 16 | A:ARG 302[ NH1] | 2.47 | B:GLU 72[ OE1] | 16 | A:ARG 299[ NH2] | 2.51 | B:GLU 75[ OE2] |
| 17 | A:ARG 299[ NH2] | 2.51 | B:GLU 75[ OE2] | 17 | A:ARG 299[ NH1] | 3.27 | B:ASP 79[ OD1] |
| 18 | A:ARG 303[ NH1] | 3.31 | B:ASP 79[ O ] | 18 | A:ARG 299[ NE ] | 2.98 | B:ASP 79[ OD2] |
| 19 | A:GLN 292[ NE2] | 3.77 | B:ASP 80[ OD1] | 19 | A:ARG 299[ NH1] | 2.85 | B:ASP 79[ OD2] |
| 20 | A:ARG 303[ NH1] | 3.54 | B:ASP 80[ OD1] | 20 | A:ARG 299[ NH2] | 3.88 | B:ASP 79[ OD2] |
| 21 | A:ARG 303[ NH1] | 2.79 | B:ASP 83[ OD2] | 21 | A:LYS 307[ NZ ] | 3.62 | B:ASP 80[ OD1] |
| 22 | A:ARG 303[ NH2] | 2.61 | B:ASP 83[ OD2] | 22 | A:ARG 303[ NH1] | 3.54 | B:ASP 80[ OD1] |
| 23 | A:THR 288[ OG1] | 2.97 | B:TYR 88[ OH ] | 23 | A:LYS 307[ NZ ] | 2.89 | B:ASP 80[ OD2] |
| 24 | A:GLY 106[ N ] | 2.90 | B:THR 89[ OG1] | 24 | A:ARG 303[ NH1] | 2.79 | B:ASP 83[ OD2] |
| 25 | A:GLU 287[ OE2] | 2.54 | B:TRP 22[ NE1] | 25 | A:ARG 303[ NH2] | 2.61 | B:ASP 83[ OD2] |
| 26 | A:GLY 106[ O ] | 3.58 | B:ARG 28[ NH2] |  |  |  |  |
| 27 | A:THR 288[ O ] | 3.01 | B:ARG 87[ NH1] |  |  |  |  |
| 28 | A:GLU 287[ O ] | 3.50 | B:ARG 87[ NH2] |  |  |  |  |

**Supplementary Table 6.**
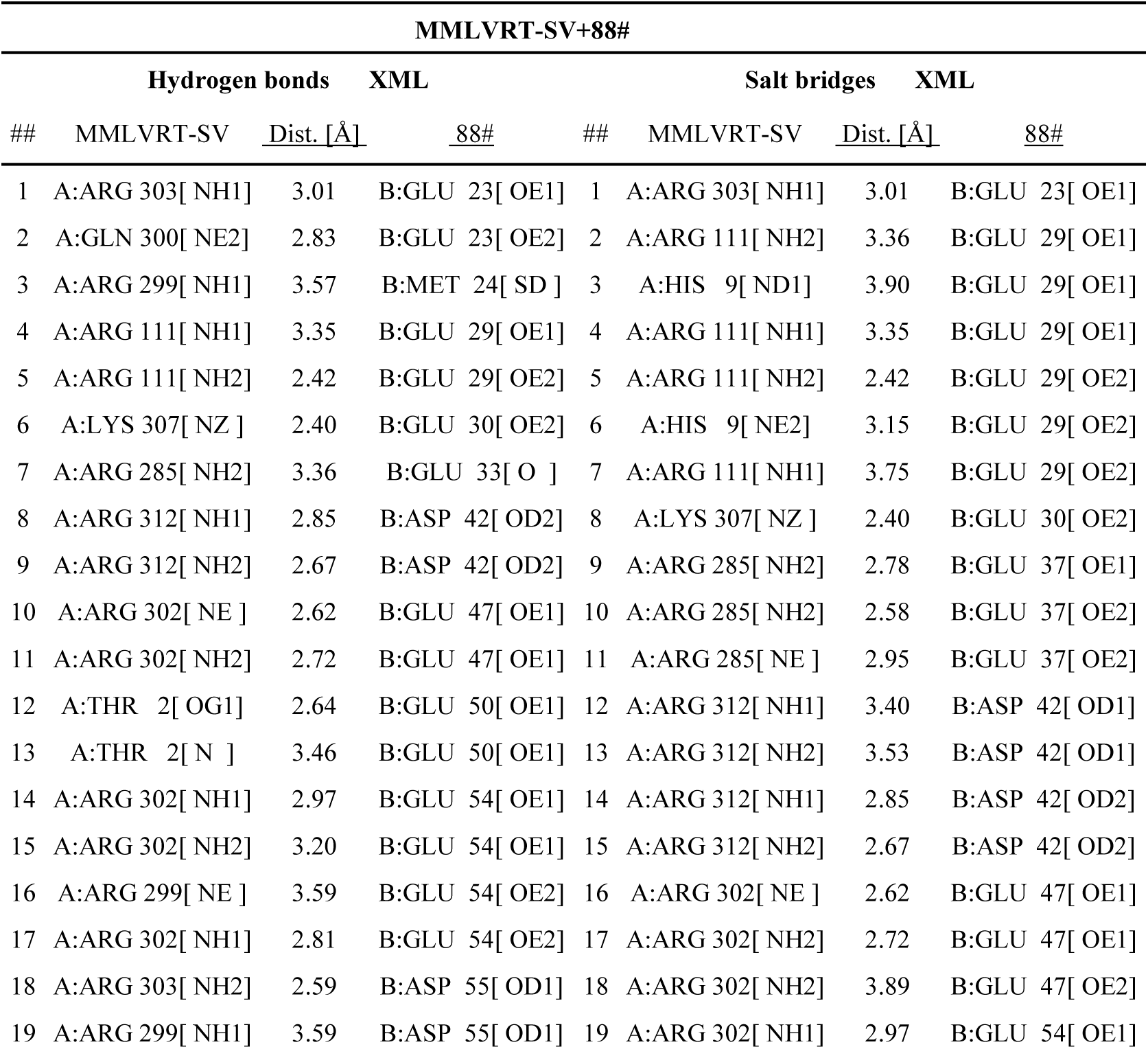
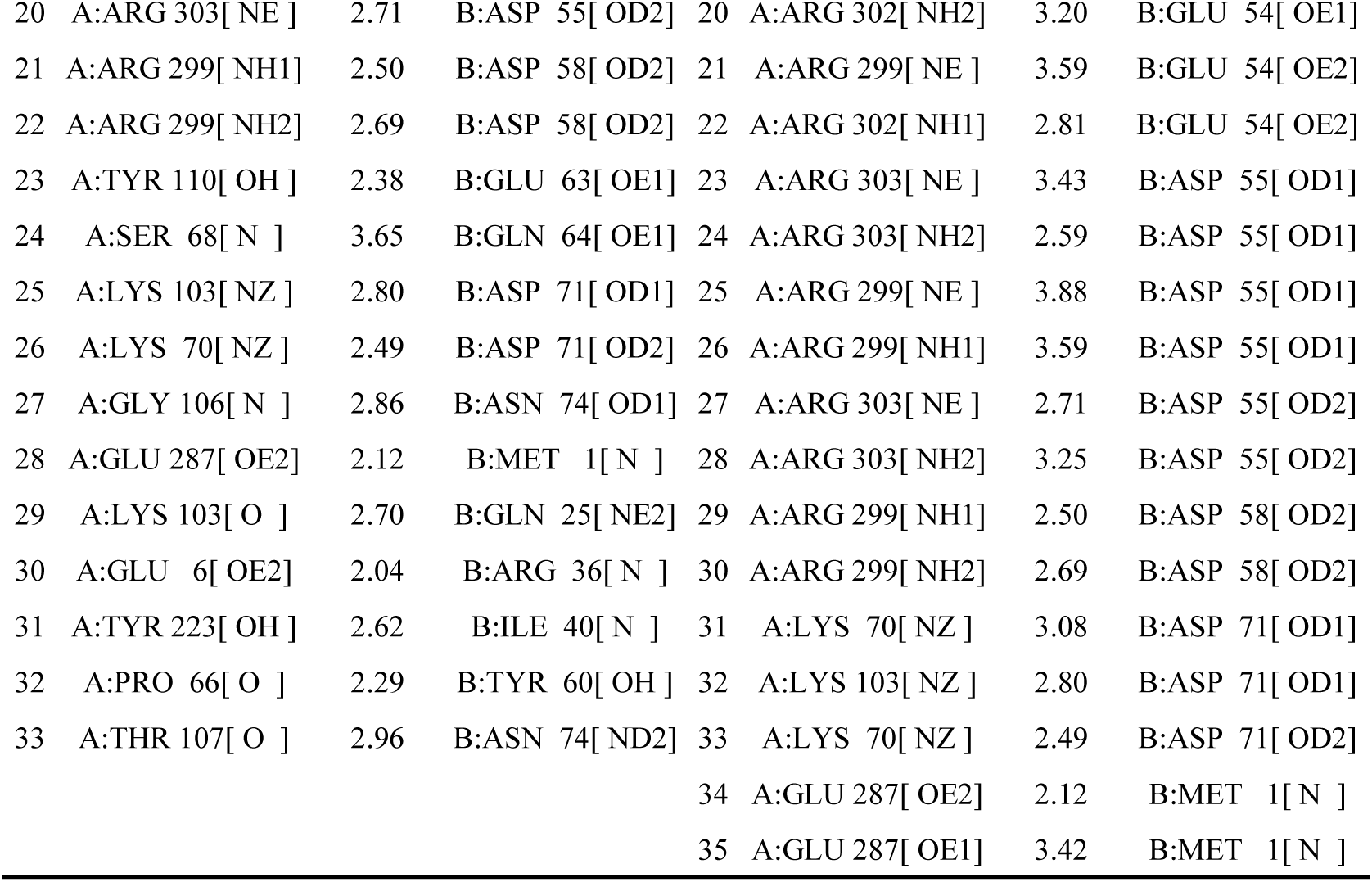
Key Residues and Interaction Statistics between Binder88# and MMLVRT-SV.

**Supplementary Table 7.**
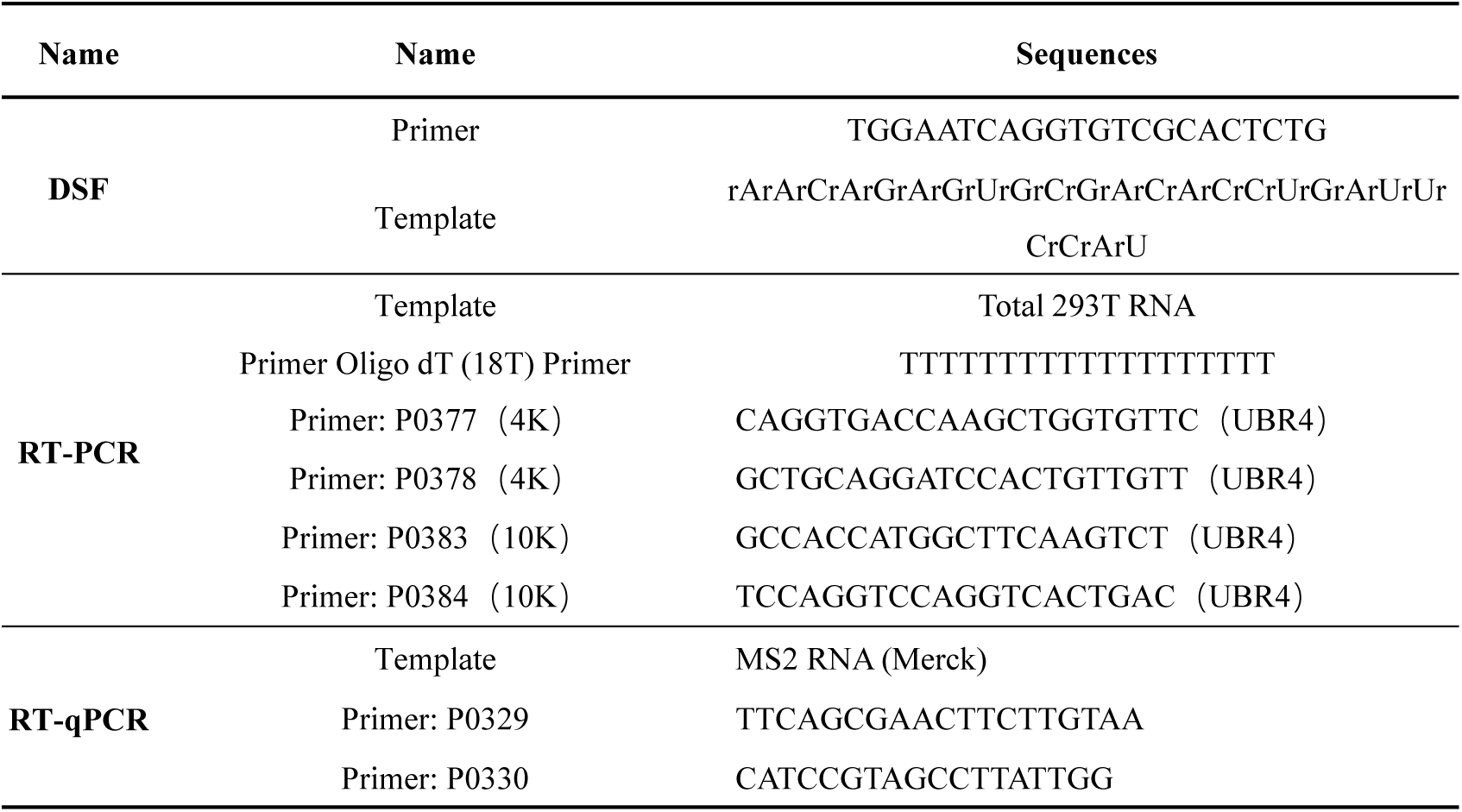
Information of Primer and Template.

**Supplementary Table 8.**
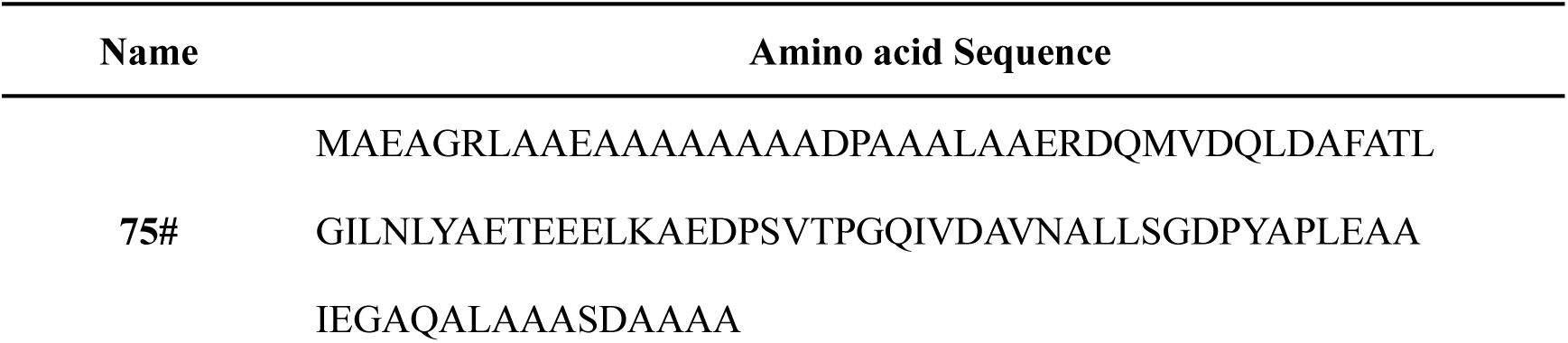
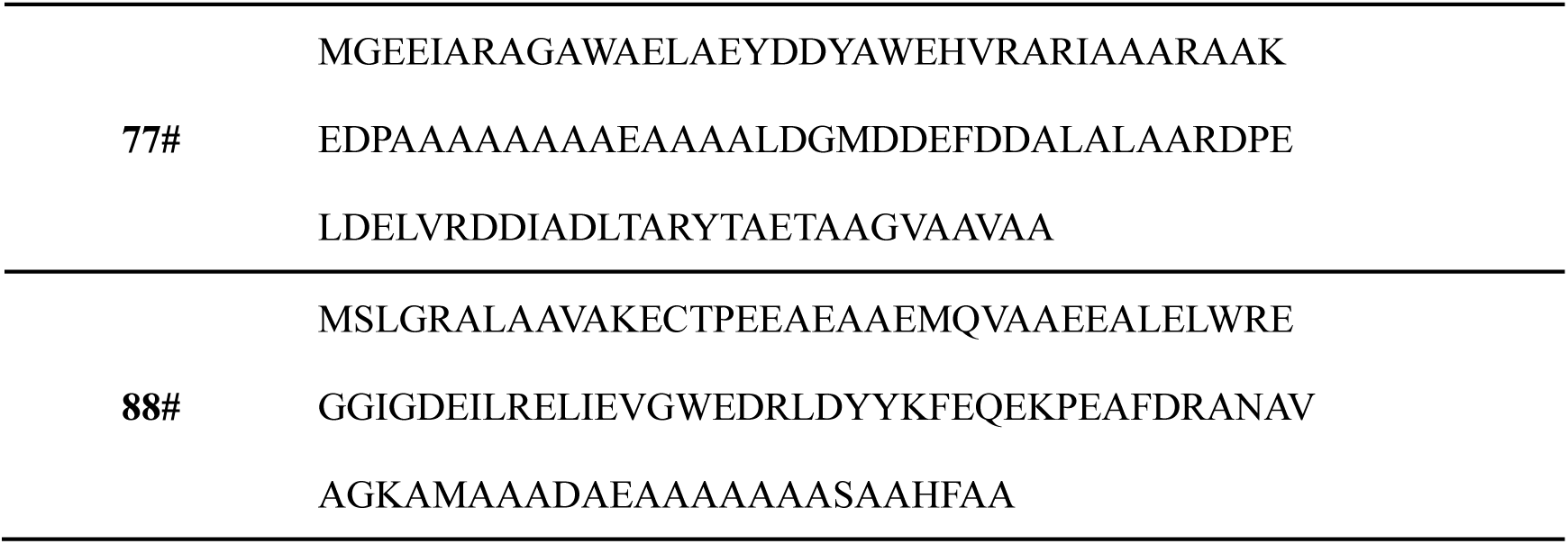
Amino acid Sequence of Binders.

## References

1. Das, D., Georgiadis, M.M. The Crystal Structure of the Monomeric Reverse Transcriptase from Moloney Murine Leukemia Virus. Structure 2004, 12, 819–829, doi:10.1016/j.str.2004.02.032.

2. Najmudin, S., Coté, M.L., Sun, D., Yohannan, S., Montano, S.P., Gu, J., Georgiadis, M.M. Crystal Structures of an N-Terminal Fragment from Moloney Murine Leukemia Virus Reverse Transcriptase Complexed with Nucleic Acid: Functional Implications for Template-Primer Binding to the Fingers Domain 1 1Edited by D. C. Rees. J. Mol. Biol. 2000, 296, 613–632, doi:10.1006/jmbi.1999.3477.

3. Grünewald, J., Miller, B.R., Szalay, R.N., Cabeceiras, P.K., Woodilla, C.J., Holtz, E.J.B., Petri, K., Joung, J.K. Engineered CRISPR Prime Editors with Compact, Untethered Reverse Transcriptases. Nat. Biotechnol. 2023, 41, 337–343, doi:10.1038/s41587-022-01473-1.

4. Shuto, Y., Nakagawa, R., Zhu, S., Hoki, M., Omura, S.N., Hirano, H., Itoh, Y., Zhang, F., Nureki, O. Structural Basis for pegRNA-Guided Reverse Transcription by a Prime Editor. Nature 2024, 631, 224–231, doi:10.1038/s41586-024-07497-8.

5. Cerda, A., Rivera, M., Armijo, G., Ibarra-Henriquez, C., Reyes, J., Blázquez-Sánchez, P., Avilés, J., Arce, A., Seguel, A., Brown, A.J., et al. An Open One-Step RT-qPCR for SARS-CoV-2 Detection 2021.

6. Graham, T.G.W., Dugast-Darzacq, C., Dailey, G.M., Darzacq, X., Tjian, R. Simple, Inexpensive RNA Isolation and One-Step RT-qPCR Methods for SARS-CoV-2 Detection and General Use. Curr. Protoc. 2021, 1, e130, doi:10.1002/cpz1.130.

7. Konishi, A., Ma, X., Yasukawa, K. Stabilization of Moloney Murine Leukemia Virus Reverse Transcriptase by Site-Directed Mutagenesis of Surface Residue Val433. Biosci. Biotechnol. Biochem. 2014, 78, 75–78, doi:10.1080/09168451.2014.877186.

8. Ohtsubo, Y., Nagata, Y., Tsuda, M. Compounds That Enhance the Tailing Activity of Moloney Murine Leukemia Virus Reverse Transcriptase. Sci. Rep. 2017, 7, 6520, doi:10.1038/s41598-017-04765-8.

9. Yang, Y., Zhang, J., Li, Z., Qi, H. Enhancing Thermostability of Moloney Murine Leukemia Virus Reverse Transcriptase through Greedy Combination of Multiple Mutant Residues. Bioresour. Bioprocess. 2025, 12, 12, doi:10.1186/s40643-025-00845-0.

10. Nishimura, K., Yokokawa, K., Hisayoshi, T., Fukatsu, K., Kuze, I., Konishi, A., Mikami, B., Kojima, K., Yasukawa, K. Preparation and Characterization of the RNase H Domain of Moloney Murine Leukemia Virus Reverse Transcriptase. Protein Expr. Purif. 2015, 113, 44–50, doi:10.1016/j.pep.2015.04.012.

11. Simanjuntak, G.M., Fibriani, A., Fananda, A.A., Yamahoki, N. Development of Moloney Murine Leukemia Virus ReverseTranscriptase Fused with Archaeal DNA-Binding Protein Sis7a. Recent Pat. Biotechnol. 2024, 18, 71–83, doi:10.2174/1872208317666230403104302.

12. Okano, H., Baba, M., Yamasaki, T., Hidese, R., Fujiwara, S., Yanagihara, I., Ujiiye, T., Hayashi, T., Kojima, K., Takita, T., et al. High Sensitive One-Step RT-PCR Using MMLV Reverse Transcriptase, DNA Polymerase with Reverse Transcriptase Activity, and DNA/RNA Helicase. Biochem. Biophys. Res. Commun. 2017, 487, 128–133, doi:10.1016/j.bbrc.2017.04.030.

13. Vanni, C., Clemente, F., Paoli, P., Morrone, A., Matassini, C., Goti, A., Cardona, F. 3,4,5-Trihydroxypiperidine Based Multivalent Glucocerebrosidase (GCase) Enhancers. ChemBioChem 2022, 23, e202200077, doi:10.1002/cbic.202200077.

14. Strandback, E., Lienhart, W., Hromic-Jahjefendic, A., Bourgeois, B., Högler, A., Waltenstorfer, D., Winkler, A., Zangger, K., Madl, T., Gruber, K., et al. A Small Molecule Chaperone Rescues the Stability and Activity of a Cancer-associated Variant of NAD(P)H:Quinone Oxidoreductase 1 *in Vitro*. FEBS Lett. 2020, 594, 424–438, doi:10.1002/1873-3468.13636.

15. Morey, T.M., Winick-Ng, W., Seah, C., Rylett, R.J. Chaperone-Mediated Regulation of Choline Acetyltransferase Protein Stability and Activity by HSC/HSP70, HSP90, and P97/VCP. Front. Mol. Neurosci. 2017, *10*, 415, doi:10.3389/fnmol.2017.00415.

16. Watson, J.L., Juergens, D., Bennett, N.R., Trippe, B.L., Yim, J., Eisenach, H.E., Ahern, W., Borst, A.J., Ragotte, R.J., Milles, L.F., et al. De Novo Design of Protein Structure and Function with RFdiffusion. Nature 2023, 620, 1089–1100, doi:10.1038/s41586-023-06415-8.

17. Lisanza, S.L., Gershon, J.M., Tipps, S.W.K., Sims, J.N., Arnoldt, L., Hendel, S.J., Simma, M.K., Liu, G., Yase, M., Wu, H., et al. Multistate and Functional Protein Design Using RoseTTAFold Sequence Space Diffusion. Nat. Biotechnol. 2024, doi:10.1038/s41587-024-02395-w.

18. Winnifrith, A., Outeiral, C., Hie, B.L. Generative Artificial Intelligence for de Novo Protein Design. Curr. Opin. Struct. Biol. 2024, 86, 102794, doi:10.1016/j.sbi.2024.102794.

19. Bennett, N.R., Coventry, B., Goreshnik, I., Huang, B., Allen, A., Vafeados, D., Peng, Y.P., Dauparas, J., Baek, M., Stewart, L., et al. Improving de Novo Protein Binder Design with Deep Learning. Nat. Commun. 2023, 14, 2625, doi:10.1038/s41467-023-38328-5.

20. Liu, H. Designing de Novo D-Protein Binders. Cell Res. 2024, 34, 820–821, doi:10.1038/s41422-024-01029-9.

